# Evolution of pathogen dormancy in fluctuating environments

**DOI:** 10.64898/2026.08.22.746398

**Authors:** Van Hai Khong, Philippe Carmona, Sylvain Gandon

**Affiliations:** Faculty of Mathematics and Informatics, Hanoi University of Science and Technology; Laboratoire de Mathématiques Jean Leray, Université de Nantes, Nantes, France; CEFE, CNRS, Univ Montpellier, EPHE, IRD, Montpellier, France

**Keywords:** dormancy, reactivation, plasticity, seasonality

## Abstract

Dormancy is a widespread life-history strategy that enables organisms to persist through periods of adverse environmental conditions. Despite its prevalence, the evolutionary forces shaping dormancy and the timing of reactivation remain poorly understood, particularly in pathogens facing predictable environmental fluctuations. Here, we investigate how seasonal variation can drive the joint evolution of pathogen dormancy and reactivation, and whether these traits are favoured to evolve as fixed or plastic strategies. Using a theoretical model of vector-borne disease transmission, we show when seasonality can promote plasticity in dormancy and reactivation. The optimal timing of transitions between active and dormant states depends critically on the environmental cues available to pathogens and on their reliability for predicting future transmission opportunities. Although motivated by the biology of relapsing malaria parasites, our results provide a general framework for understanding the evolution of dormancy as an adaptive response to periodic environmental fluctuations across diverse pathogen systems.

## 1 Introduction

Many different species produce offspring that stop their development and stay in a dormant stage, sometimes for a very long time [12]. This ability to produce dormant stages is thought to be an adaptation to the temporal variability of the environment [21, 34, 36]. For instance, many microbes have the ability to delay growth to cope with periods where growth conditions become unfavorable [3, 6]. Here we examine the evolution of dormancy in a vector-borne pathogen under the influence of seasonal variations of the abundance of vectors. We consider a scenario where the pathogen can potentially switch between two host-exploitation strategies: (i) the pathogen can replicate actively in the host in the hope to be picked up by a vector and transmitted to a new host; (ii) the pathogen can lay dormant in the host tissue and avoid the risk of being targeted by the host immune system, but reduce its ability to infect a new host. These two alternative strategies are meant to describe different developmental stages. For instance, after infecting a human host, the sporozoites of the malaria parasite *Plasmodium vivax* can either develop as a merozoite and replicate asexually in the bloodstream of its host, or lay dormant in the liver as an hypnozoite. Hypnozoites can remain dormant for several months but they can eventually relapse and differentiate into merozoites. It is interesting to note that different isolates of *P. vivax* relapse at different rates which may indicate the existence of some heritable variation on this trait [38, 39]. In *P. vivax*, the merozoites cannot switch back to a dormant state but some avian malaria species are known to have the ability to switch back and forth between a dormant and a non-dormant state [31]. Other malaria species (e.g. *P. falciparum*) do not have the ability to produce hypnozoites which also suggest that dormancy may not always be adaptive. Finally, we also want to note that some malaria parasites have the ability to adopt plastic strategies and to switch between different developemental stages in response to a change of the environment [24, 28, 32, 40]. For instance, mosquito bites can trigger a higher investment into pathogen transmission in the avian malaria parasite *P. relictum* [8]. The observed diversity of life-history strategies among malaria parasites may result from adaptation to different levels of seasonality. Yet, the analysis of the the evolutionary consequences of seasonality for vector-borne pathogens remain only partially understood. Earlier models have explored the effect of seasonality on the evolution of fixed or plastic relapsing strategies [8, 37]. But a general theoretical framework accounting for the class-structure of these models for predicting how seasonality shapes the evolution of vector-borne pathogens is still lacking.

To address this gap we develop a general theoretical framework that integrates epidemiology and evolution while explicitly accounting for both class-structure and seasonal fluctuations of the environment. First, we study the influence of seasonality on epidemiological dynamics and pathogen persistence. Second, we derive general selection gradients on pathogen dormancy and reactivation under different regimes of seasonality. Third, we use these selection gradients to study the joint evolution of dormancy and reactivation in a constant or in a temporally variable environment. We contrast different scenarios where (i) dormancy and reactivation are fixed traits or (ii) the investment in dormancy and reactivation is conditional on different environmental cues. Finally, we discuss the implications of our theoretical results for the evolution of complex life-history strategies of pathogens living in temporally variable environments.

## 2 The model

Three organisms interact in the life-cycle of malaria parasites: the vertebrate host (noted with the subscript *H*), the mosquito vector (noted with the subscript *V*), and the malaria parasite which may be infectious (noted *I*) or dormant (noted *D*). This yields an epidemiological model with 6 different compartments (**Figure 1**): the vertebrate host may be susceptible, infectious, infected by a dormant stage or recovered (*S*_*H*_, *I*_*H*_, *D*_*H*_ or *R*_*H*_) ; the mosquito vector may be susceptible or infectious (*S*_*V*_ or *I*_*V*_). We assume that when the malaria parasite is in a dormant stage, it is less visible by the immune system of the vertebrate host and the parasite is cleared at a lower rate than in an infectious stage (*γ*_*D*_ *< γ*_*I*_). On the other hand, a dormant infection is assumed to have a lower infectivity and the parameter *c*_*D*_ ∈ [0, 1] measures the cost of dormancy on malaria transmission. When *c*_*D*_ = 0, the compartment *D*_*H*_ is as infectious as the compartment *I*_*H*_. In contrast, if *c*_*D*_ = 1, the compartment *D*_*H*_ is not infectious and the pathogen has to reactivate and go back to the compartment *I*_*H*_ to transmit to new hosts. This yields the following set of ordinary differential equations:

**Figure 1:**
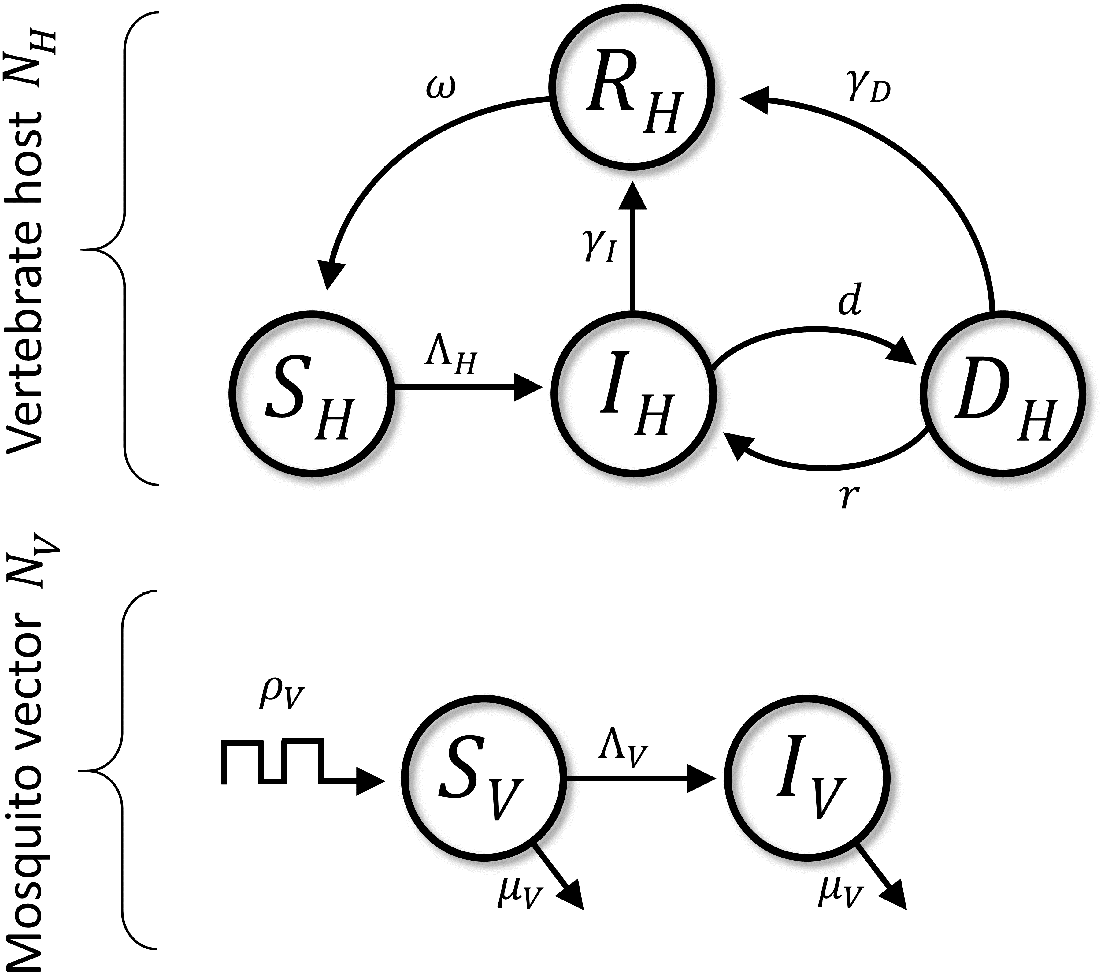
Schematic presentation of our model of the malaria life cyle. The model tracks the dynamics of four states of the vertebrate host and the two states of the mosquito vector (see also **Table 1**). Transitions from the susceptible states to the infectious states are driven by the forces of infection Λ_*H*_ (*t*) = *β*_*V*_ *I*_*V*_ (*t*)*S*_*H*_ (*t*) and Λ_*V*_ = *β*_*H*_ (*I*_*H*_ (*t*) + (1 − *c*_*D*_)*D*_*H*_ (*t*)) for the host and the vector, respectively. Because the death rate compensates exactly the birth rate of the host the total population size of the host population is assumed to be constant: *N*_*H*_ = *S*_*H*_ (*t*) + *I*_*H*_ (*t*) + *D*_*H*_ (*t*) + *R*_*H*_ (*t*) = 1. In contrast, since the reproduction of the vector is seasonal, its total density 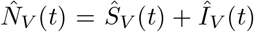 fluctuates periodically according to (2.3).

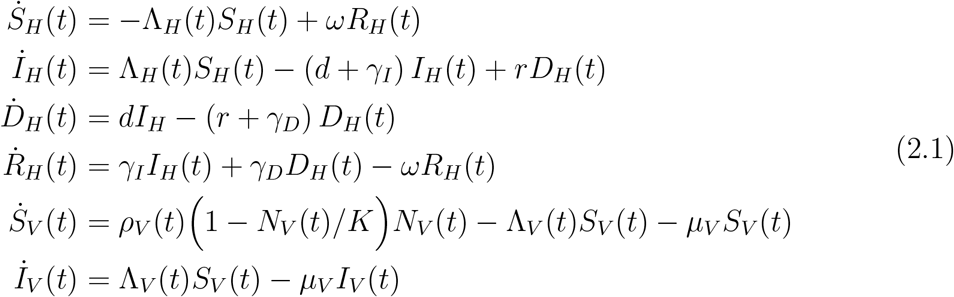

where Λ_*H*_(*t*) = *β*_*V*_ *I*_*V*_ (*t*) and Λ_*V*_ (*t*) = *β*_*H*_ (*I*_*H*_(*t*) + (1 − *c*_*D*_)*D*_*H*_(*t*)) refer to the forces of infection on the vertebrate host and on the mosquito vector, respectively. The density of the vertebrate host population is assumed to be constant so that *N*_*H*_ = *I*_*H*_(*t*) + *S*_*H*_(*t*) + *D*_*H*_(*t*) + *R*_*H*_(*t*). Without loss of generality, we will assume *N*_*H*_ = 1. Other parameters of the model are described in **Table 1**.

**Table 1:** Parameters and variables of the models. In a seasonal environment we assume that the growth rate *ρ*_*V*_ (*t*) of the vector population varies between 0 in the “low transmission season” and *ρ* = 1 *day*^*−*1^ in the “high transmission season”. The acquired immunity to malaria is assumed to be short-lived and we thus assume *ω* = 1 *day*^*−*1^. All other parameter values are obtained from [37], where *h* and *m* refer to the units used to measure the densities of vertebrate hosts and the densities of mosquito vectors, respectively. As explained in the main text, without loss of generality we assume *N*_*H*_ = 1*h*.

| Symbol | Description | Default value |
| --- | --- | --- |
| $\beta_V$ | per capita rate of transmission from $I_V$ to $S_H$ | $0.0316 m^{-1} \cdot \text{day}^{-1}$ |
| $\beta_H$ | per capita rate of transmission from $I_H$ to $S_V$ | $0.0483 h^{-1} \cdot \text{day}^{-1}$ |
| $\rho$ | maximal growth rate of the vector population | $3 \text{ day}^{-1}$ |
| $K$ | carrying capacity of the vector population | $10 m$ |
| $\mu_V$ | death rate of the vector | $0.1 \text{ day}^{-1}$ |
| $\gamma_I$ | rate of clearance of $I_H$ hosts | $1/60 \text{ day}^{-1}$ |
| $\gamma_D$ | rate of clearance of $I_D$ hosts | $1/223 \text{ day}^{-1}$ |
| $\omega$ | rate of waning of immunity | $1 \text{ day}^{-1}$ |
| $T$ | period of the seasonal fluctuations | $365 \text{ days}$ |
| $\sigma$ | duration of the low-transmission season (“winter”) | $0.4$ |
| $d$ | rate of dormancy | evolving trait |
| $r$ | rate of reactivation | evolving trait |
| $c_D$ | cost of dormancy on malaria transmission | variable |
| $t_s^d$ | starting time of dormant period | evolving trait |
| $d_{\text{on}}$ | dormancy when it is switched on | $5/365 \text{ day}^{-1}$ |
| $t_e^d$ | ending time of dormant period | evolving trait |
| $d_{\text{off}}$ | dormancy when it is switched off | $0 \text{ day}^{-1}$ |
| $t_s^r$ | starting time of reactivation | evolving trait |
| $r_{\text{on}}$ | reactivation when it is switched on | $5/365 \text{ day}^{-1}$ |
| $t_e^r$ | ending time of reactivation | evolving trait |
| $r_{\text{off}}$ | reactivation when it is switched off | $0 \text{ day}^{-1}$ |

Crucially, we assume that the density of the vector population may fluctuate periodically in time because of seasonality (where *T* is the period of the fluctuation). The dynamics of the density *N*_*V*_ (*t*) = *S*_*V*_ (*t*) + *I*_*V*_ (*t*) of the vector population is driven by the balance between the fluctuations of the time-varying reproduction rate *ρ*_*V*_ (*t*) and the constant death rate *µ*_*V*_ which yields:

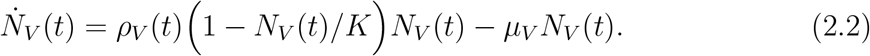

We assume that *ρ*_*V*_ (*t*) = 0 when the environment is not favorable for the vector (i.e. winter), and *ρ*_*V*_ (*t*) = *ρ* when the vector can reproduce (i.e. summer). In addition, we assume that the parameter *σ* measures the relative duration of the winter (i.e. the proportion of the year). Seasonality leads to periodic dynamics and we can show that the vector population is expecetd to sit on the following periodic attractor 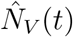, where the refers to the attractor (see Supplementary Information section S1):

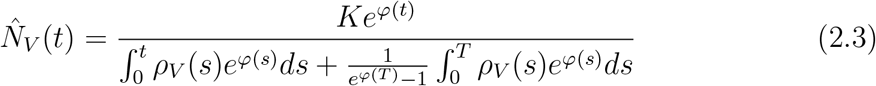

where 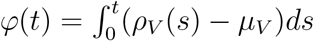

## 3 Results

Before studying the evolution of the pathogen we must verify that the pathogen can persist in the host population. In other words, we have to verify that the basic reproduction ratio of the pathogen satisfies the condition *R*_0_ *>* 1. Yet, the derivation of the basic reproduction ratio can be difficult in a periodically fluctuating environment. We extend the perturbation analysis used by [19] to derive an approximation for *λ*, the asymptotic growth rate of the epidemic in the initial phase of the epidemic when it is computed over on period of the fluctuation. Crucially, both *R*_0_ and *λ* can be used to analyse the persistence of the pathogen population because *R*_0_ −1 and *λ* −1 have the same sign [19]. We analyse in the Supplementary Information S1 the persistence of the pathogen population in constant or seasonal environments. In particular, if we perturb the equilibrium near the point *λ* = 1 for small values of *σ* (the parameter that measures the duration of the winter) we can write *λ* = *λ*_0_ + *σλ*_1_ + *o*(*σ*) where *λ*_0_ refers to the case without seasonality (i.e. *σ* = 0), and *λ*_1_ refers to the first-order effects of seasonality on pathogen persistence. Not surprisingly, we find that *λ*_1_ *<* 0 which means that seasonality has a negative effect on pathogen persistence (**Figure S1**). This effect is due to the overall drop in the density of vector (and thus in transmission opportunities) when duration of the winter season increases (see proposition S1.2 in the Supplementary Information). Next, we focus on scenarios where *λ >* 1 and we study the phenotypic evolution of the pathogen.

### 3.1 The selection gradient

Following Lion and Gandon [23] we derive the selection gradient associated with a specific mutation affecting the phenotype *z* of the pathogen (*z* = *d* or *r* to refer to dormancy or reactivation, respectively) in a periodically fluctuating environment. We assume that the phenotype *r*_*m*_ of the mutant is close to the resident phenotype so that *z*_*m*_ = *z*_*w*_ + *ϵ*, with *ϵ* small. This allows us to obtain a weak selection approximation of the selection gradient to determine the fate of the mutant (invasion or extinction) and, ultimately, to identify evolutionary stable strategies.

To follow the dynamics of the mutant we track the dynamics of the vector 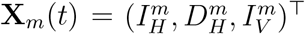, with:

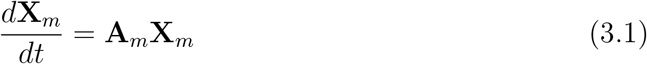

with

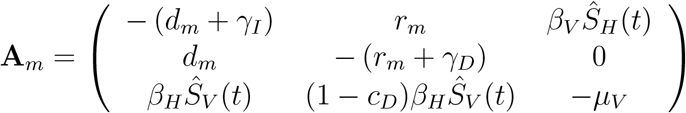

The coefficients *a*_*jk*_ of the matrix **A**_*m*_ are the transition rates from an infection in state *k* to a new infection in state *j*. Note that some of these coefficients depend on *Ŝ*_*H*_ (*t*) and *Ŝ*_*V*_ (*t*), the fluctuating densities of the host and the vector when the system sits on the periodic attractor. These fluctuations affect both the quantity and the quality of the three different types of infected compartments(*I*_*H*_, *D*_*H*_ and *I*_*V*_). The selection gradient on the mutant at time *t* is (see Lion and Gandon [23]):

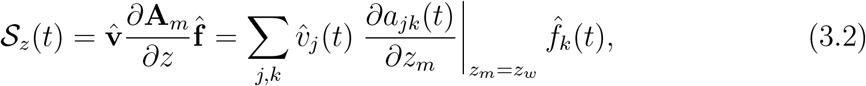

where *f*_*k*_ and *v*_*j*_ refer to the frequency and the individual reproductive value of pathogens in the infected compartments *k, j* ∈L{*I*_*H*_, *D*_*H*_, *I*_*V*_}, respectively. Note that these quantities have been normalized so that *Σ*_*k*_ *f*_*k*_ = 1 and *Σ*_*j*_ *f*_*j*_*v*_*j*_ = 1.
The selection gradient on dormancy and reactivation can thus be obtained from averaging the selection coefficient over one period of the fluctuation of the resident attractor which yields:

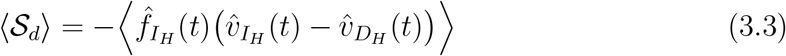

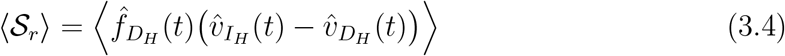

where we use the notation 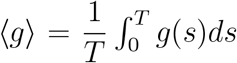 to indicate the average over one period of the fluctuation. To determine the direction of phenotypic evolution we thus need to compute the class frequencies and the reproductive values of the different infection states when the system sits on the periodic attractor (see Supplementary Information S2). Next, we use the above selection gradients to study the evolution of the pathogen under different ecological scenarios when the pathogen adopts fixed dormancy and reactivation strategies (i.e. without plasticity).

### 3.2 Pathogen evolution without plasticity

#### Constant environment

Let us first focus on the scenario where the environment does not fluctuate with time. In other words, we assume there is no seasonality and *σ* = 0. In this case we can use (3.3) and (3.4) to show:

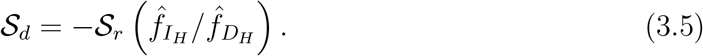

Hence, in a constant environment, selection for higher rates of dormancy always yield selection for lower rates of reactivation. In other words, we expect the two traits to evolve in opposite directions. Besides, in a constant environment, we can compute the ratio of reproductive values (Supplementary Information S2.1) which yields:

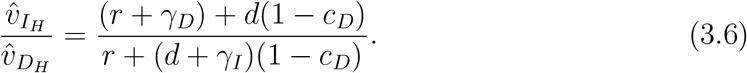

This implies that selection for dormancy (reactivation) is positive (negative) whenever 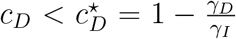. Indeed, this threshold measures the probability that the infection is cleared by the host immune system before the death of the host. When the cost of dormancy on transmission drops below this threshold value, the reproductive value of the dormant stage becomes higher than the reproductive value of an infectious stage (because the dormant stage is protected from clearance and can still transmit). The dashed red line in **Figure 2** illustrates the effect of the cost of dormancy *c*_*D*_ on the evolution of the pathogen. In a constant environment, dormancy is either maximized or minimized, depending on the cost of dormancy relative to 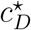 (and vice versa for reactivation). Next we analyse how the seasonality of the environment may affect the evolution of the pathogen.

**Figure 2:**
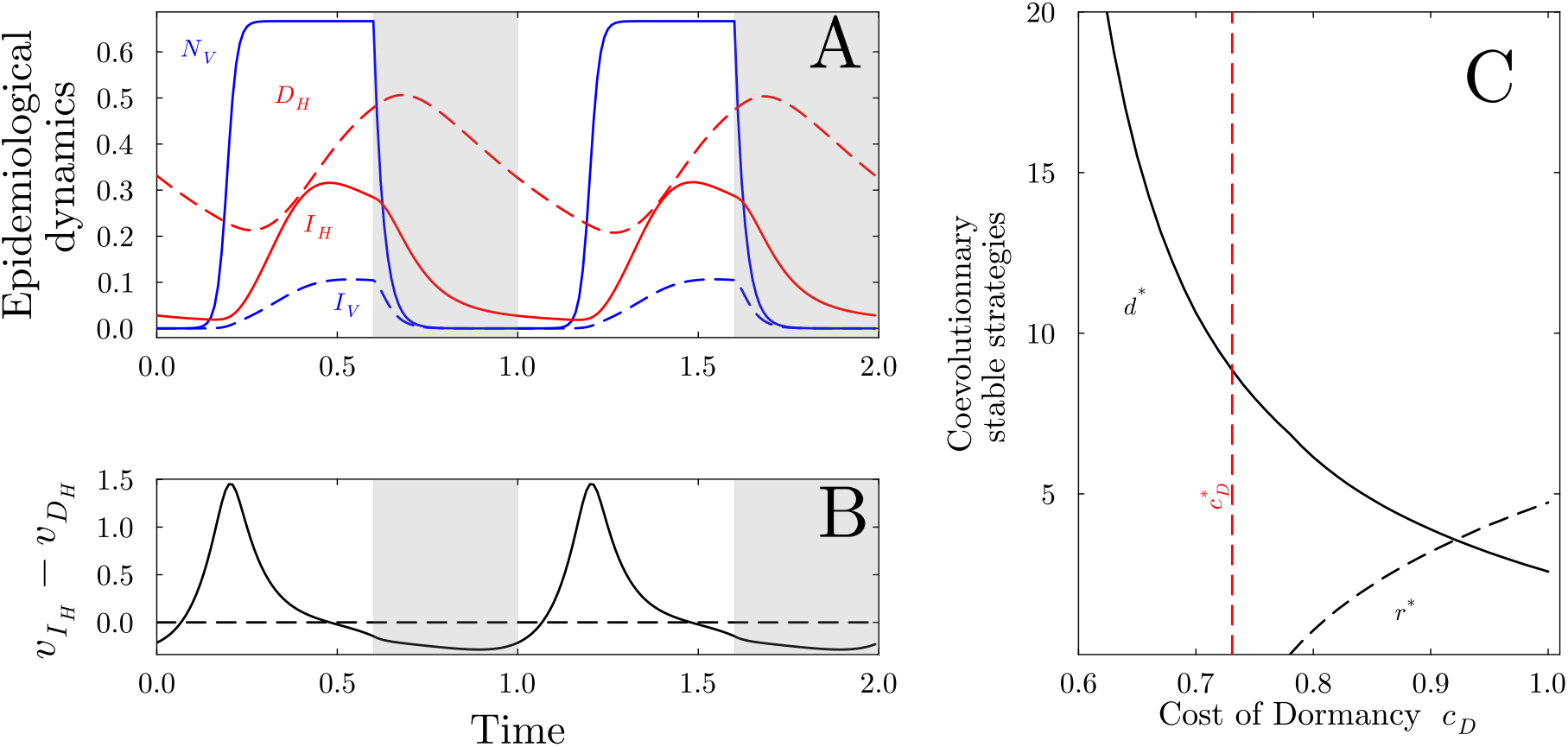
Coevolution of dormancy and reactivation in a seasonal environment. Panel (A) shows the fluctuations of the densities of different types of vertebrate hosts (*I*_*H*_ : dashed full line, *D*_*H*_ : dashed red line) and mosquito vectors (*I*_*V*_ : dashed blue line, *N*_*V*_ : full blue line) in a seasonal environment with *σ* = 0.4 (the gray area indicate the winter season, when mosquito stop reproducing). Panel (B) illustrates the consequences of the epidemiological fluctuations on the dynamics of the difference between the reproductive values of *I*_*H*_ and *D*_*H*_ which drives the selection for dormancy and reactivation. Panel (C) shows the coevolutionary stable strategies of dormancy and reactivation, *d*^*∗*^ and *r*^*∗*^ for different values of the cost of dormancy *c*_*D*_. The dashed vertical line in red gives the value of 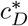, the threshold value of the cost driving the coevolution of dormancy and reactivation in the non-seasonal environment. In panels (A) and (B) we use *c*_*D*_ = 0.8 and we stand at the coESS *d*^*∗*^ = 6.15*/*365 *day*^*−*1^ and *r*^*∗*^ = 0.75*/*365 *day*^*−*1^. Other parameter values are given in Table 1. In the **bottom left**, for the same parameters, we observe the difference of reproductive values 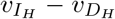 : when 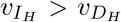 reactivation is favored, when 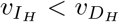 dormancy is favored. Other default parameters are given in **Table 1**.

#### Seasonal environment

Seasonality induces periodic fluctuations of the densities of mosquito vectors (**Figure 2A**). These fluctuations affect the incidence of the disease and, consequently, the selection gradient for dormancy and reactivation (see equations 3.3 and 3.4) via the dynamics of 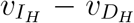 (**Figure 2B**). For instance, 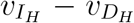 tends to be negative during the winter season (when the density of mosquitoes drops) which selects for dormancy. If we allow dormancy and reactivation to coevolve, we can look for the coevolutionary stable strategy (coESS) that verifies ⟨*S*_*d*_⟩ = ⟨*S*_*r*_⟩ = 0. In contrast with the constant environment, seasonality can promote the evolution of mixed coevolutionary strategies (*d*^*∗*^, *r*^*∗*^) (**Figure 3**). Indeed, seasonality drives fluctuations of class frequencies and reproductive values and the selection gradient depends on the temporal covariance between these quantities (see Supplementary Information). As expected, we find that the coESS depends also on the cost of dormancy *c*_*D*_. We show in **Figure 2C** that lower rates of dormancy and higher rates of relapse evolve with higher costs of dormancy. Increasing the duration *σ* of the “winter” affects the coevolution of dormancy and reactivation (**Figure 3**). When *σ* is very low, the pathogen experiences a high-transmission environment most of the year and we recover the results obtained in the constant environment: the dormancy switches “off” (and reactivation switches “on”) when 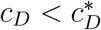. Larger values of *σ*, however, allow for intermediate coESS *d*^*∗*^ and *r*^*∗*^ for a broad range of values for the cost of dormancy.

**Figure 3:**
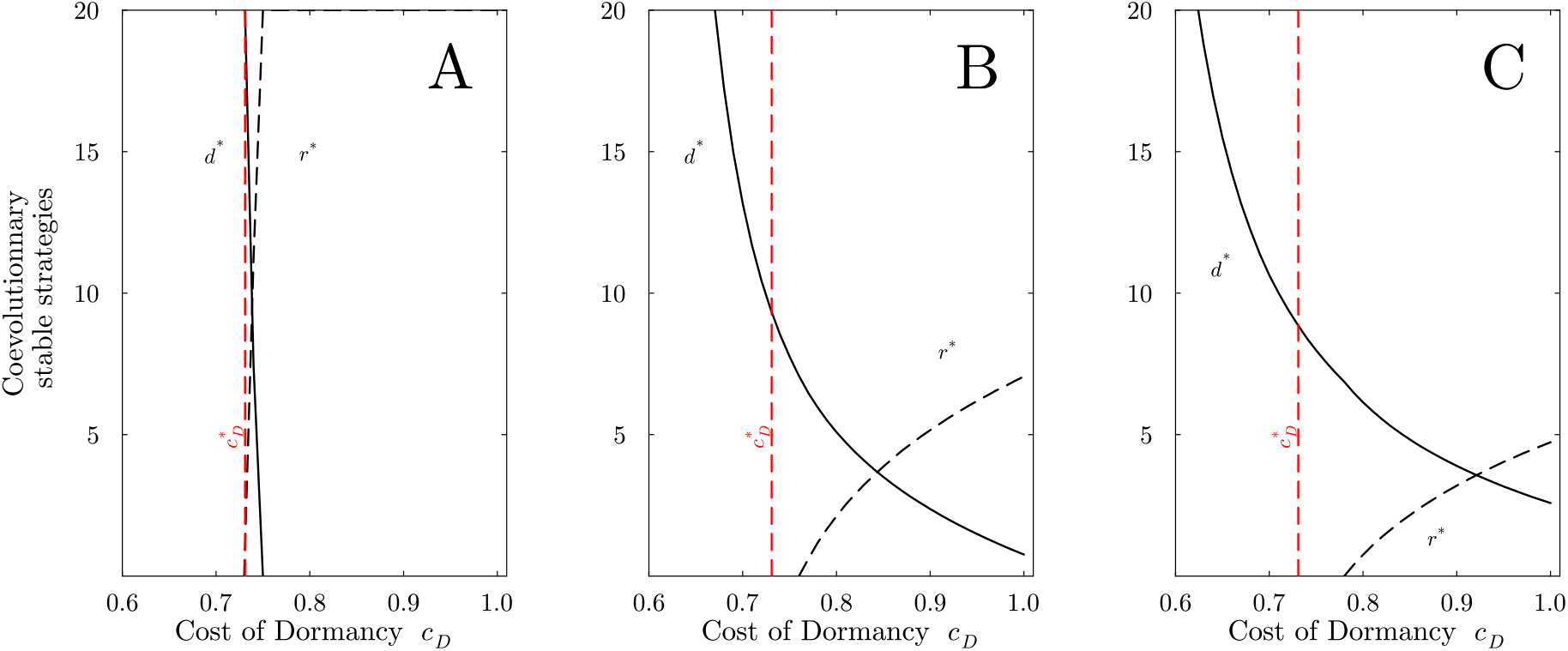
Coevolution of dormancy and reactivation for increasing levels of seasonality. We show the coevolutionary stable strategies of dormancy and reactivation, *d*^*∗*^ and *r*^*∗*^ for different values of the cost of dormancy *c*_*D*_ and for increasing levels of seasonality: (A) *σ* = 0.1, (B) *σ* = 0.3, (C) *σ* = 0.5. The dashed vertical line in red gives the value of 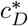, the threshold value of the cost driving the coevolution of dormancy and reactivation in the non-seasonal environment. Other default parameters are given in **Table 1**.

### 3.3 Pathogen evolution with plasticity

In the following we relax the assumption that the pathogen strategies *d* and *r* are fixed. We now consider that the pathogen can perceive the variations of some components of its environment (i.e. cues) and use these variations to modulate *d* and *r*. In other words, we consider that *d*(*t*) and *r*(*t*) can vary plastically with time in response to different cues that provide some information on the environment. We consider two different types of cues.

#### Plasticity with time

We first examine a situation where the pathogen is able to keep track of time and to switch “on” and “off” dormancy (or reactivation) at specific points in time. More specifically, we assume that (i) the pathogen invests into dormancy at rate *d*_on_ during the time interval 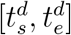 (outside this interval we assume *d*_off_ = 0), (ii) the pathogen invests into reactivation at rate *r*_on_ during the time interval 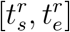 (outside this interval we assume *r*_off_ = 0). In other words, we assume:

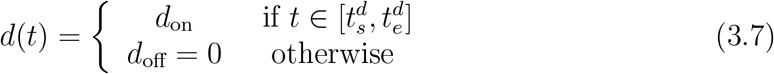

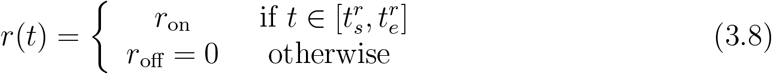

We derive the selection gradients on the switch times for dormancy and reactivation in Supplementary Information S2.2. We find that dormancy should be switched “on” when reactivation is switched “off”, and vice versa. In other words, the plastic coESS depends only on two traits 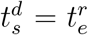 and 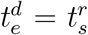. **Figures 4A-C** show the ESS plastic strategy for decreasing levels of the cost of dormancy. As expected, the pathogen switches “on” dormancy around the winter season, but the timing of dormancy period depends on the cost of dormancy. Note that the plastic ESS strategy of the pathogen switches to dormancy ahead of the winter season because the pathogen anticipates the low transmission season. Note that when the cost of dormancy is very low (i.e. the infectivity of dormant infections remain relatively high), the pathogen may never reactivate and remain in the dormant state throughout the year. Besides, even if dormancy is always “on” the pathogen spends a time 1*/d*_*on*_ in the fully infectious class in the early stage of the infection (this period of time becomes vanishingly small when *d*_*on*_ increases). The same qualitative results hold when we vary *σ*, the length of the winter season **(Figure S3-S6**). The dormancy period gets shorter when the winter season is shorter and it can even vanish completely (**Figure S3**). Yet, when the duration of the winter is long and the cost of dormancy is low, the pathogen may evolve strategies that reactivate during the short period of time where mosquito vectors are around (**Figure S6C**).

**Figure 4:**
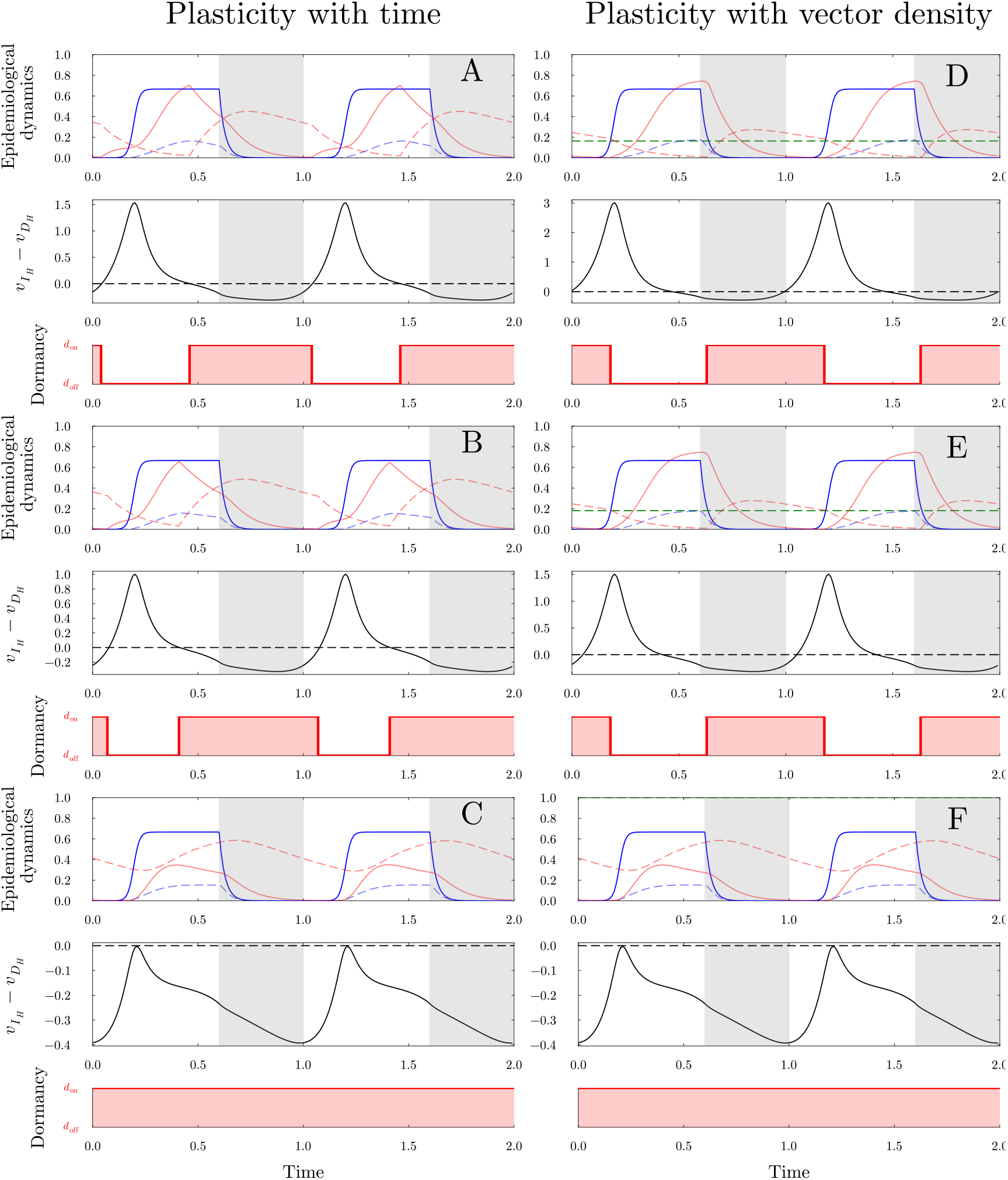
Evolution of time-varying dormancy in a seasonal environment. Each panel presents the epidemiological dynamics (same notation as in **Figure 2A**), the fluctuations of the difference between the reproductive values 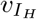 and 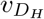 and the plastic dormancy strategy under two different assumptions. In panels (A-C), we assume that the pathogen is able to switch its strategy at any point of time during the season. In panels (D-F), we assume that the pathogen is only able to perceive time through the density of mosquitoes *N*_*V*_ (*t*). Hence, the beginning (or the end) of the winter is perceived when this density goes above (or below) a threshold value 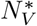 (the green dashed line). This constraint affects the timing of the switch of the evolutionary stable dormancy strategy *d*^*∗*^. We vary the cost of dormancy: *c*_*D*_ = 1 in (A) and (D), *c*_*D*_ = 0.85 in (B) and (E), *c*_*D*_ = 0.4 in (C) and (F). Other parameter values are are given in **Table 1**.

#### Plasticity with vector density

Here we examine an alternative scenario where the pathogen is unable to track the time of the year, but uses the density of vectors *N*_*V*_ (*t*) as an indirect cue to track the change of the environment [8]. Hence, the pathogen can switch “on” and “off” dormancy (or reactivation) when the vector density goes beyond or below a certain threshold value 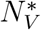 . More specifically, when 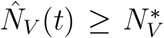 the pathogen reactivates the dormant hosts at rate *r* = *r*_on_ and dormancy is switched off (*d* = 0). In contrast, when 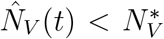 dormancy is switched “on” *d* = *d*_on_ and reactivation is switched off *r* = 0. We thus assume:

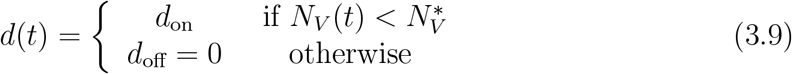

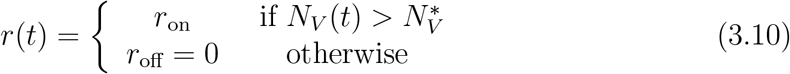

We derive the selection gradients on the switch times for dormancy and reactivation in Supplementary Information S2.2. Because the ESS depends on a single parameter, the threshold density 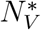, the evolution is more constrained than in the previous scenario and we obtain different switch times for the ESS strategy. In particular, we find that the investment in dormancy tends to end later after the winter because there is a lag in the population dynamics of the density of vectors (**Figure 4**). Also, the timing of the phenotypic switch is less sensitive to a variation of the cost of dormancy because the population dynamics of the vector is not affected directly by this cost. Yet, when this cost is very low, both strategies can yield a permanent investment in dormancy (**Figure 4C** and **4F**).

## 4 Discussion

We study the joint evolution of pathogen dormancy and reactivation in vector-borne pathogens in a seasonal environment characterized by a periodic fluctuations between high- and low-transmission seasons. We assume that the dormant state is less infectious (i.e. the cost of dormancy *c*_*D*_ *>* 0) but it allows the pathogen to hide from host immunity during the low-transmission season (i.e. *γ*_*D*_ *< γ*_*I*_). Our model goes beyond the analysis carried out in [8, 27] because we do not assume a separation of time scale between the processes taking place in the vertebrate and the mosquito hosts. Hence, we can explore the effect of seasonality on the evolution of a broader range of fixed or plastic phenotypic traits.

First, we focus on the analysis of the coevolution between fixed rates of dormancy and reactivation. We show that coESS strategies are governed by the cost of dormancy and the duration of the low-transmission season. In the absence of the low-transmission season, vector density is constant and the pathogen evolves a fully non-dormant strategy if the cost of dormancy is above a threshold 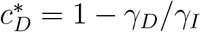, and it evolves a fully dormant strategy if the cost of dormancy is below this threshold. Hence, in low-latitude environments, where mosquito vectors are present throughout the year, differences in the cost of dormancy among different pathogens may explain the differences in dormancy among various *Plasmodium* pathogens. Yet, most habitats are subject to some level of seasonality and our model allows us to explore its effect on the evolution of dormancy. In particular, we show that in high-latitude environments, where seasonality is maximal and the low-transmission season is long, we expect the evolution of higher rates of dormancy even when the cost of dormancy is very high. This prediction is in line with the observation that isolates of *P. vivax* sampled at high latitudes tend to exhibit lower rates of reactivation (i.e. longer latency period and lower frequency of relapses) than isolates sampled at low latitudes [8, 38].

Next, we allow the pathogen to evolve plastic investment in dormancy and reactivation. In this case, we assume that the detection of a change of a specific component of the environment (an environmental cue) by the pathogen may trigger a switch in dormancy and/or reactivation. We consider two types of cues. First, we assume that the pathogen may be able to perceive seasonal changes through physiological or immunological changes of their host driven, for instance, by the variation of the photoperiod [1, 2, 10, 26, 35]. Under this scenario the pathogen can time its investment in dormancy and reactivation with the seasons. Our analysis shows that the evolutionary stable switching times depend critically on the cost of dormancy and on the duration of the low-transmission season. Interestingly, the pathogen is expected to anticipate the low-transmission season and invests into dormancy before the winter. Second, we consider another type of cue where the pathogen can only track seasonal changes indirectly. Under this scenario, we assume the pathogen is able to detect the presence of mosquitoes when they bite their host [17, 18, 18, 22, 25]. In this case, the pathogen may switch its phenotypes when the vector density reaches a threshold 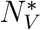 and we allow this threshold to evolve freely. Yet, this environmental cue implies that the same threshold is used to switch “on” (when the number of bites is above the threshold) or switch “off” dormancy (when the number of bites is below the threshold). This constraint affects the evolutionary stable switching times. In particular, we find that the ESS investment into dormancy is delayed in time and starts after the beginning of the winter because the fluctuations of mosquito density (the cue) always lags behind the seasonal fluctuations. The above scenario assumes that dormancy is switched “on” when reactivation is switch “off” and vice versa. We relaxed this constraint in an alternative scenario where the threshold 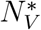 may vary between dormancy and reactivation and we find that, in this case, dormancy and reactivation are not expected to adopt bang-bang strategies where dormancy is “on” when reactivation is “off”, and vice versa (**Figure S7**). These two different scenarios illustrate how different mechanistic constraints on the perception of seasonal changes can affect the phenotypic evolution of the pathogen. It would be interesting to explore the investment in dormancy stages across a latitudinal gradient to test the validity of our predictions on the effect of *σ*, the duration of the winter season. In particular, evidence that some pathogens can anticipate seasonal changes may provide key insights on the underlying mechanism used to perceive the fluctuations of its environment.

Our numerical exploration focuses on seasonal changes with a period of one year but our model could also be used to study the evolution of pathogens exposed to shorter or longer periodic fluctuations of their environment. In particular, several pathogens have the ability to vary their investment into transmission throughout the day to coincide with the activity of their mosquito vectors [27, 29]. Several studies explored experimentally the coupling between the regulation of the circadian rhythm of the pathogens and its host has been studied experimentally [7, 15, 16]. Our model could complement this approach by studying the impact of the daily fluctuations of the activity of the vectors on the evolution of the regulation of within-host dynamics of the pathogen. Our model may also be relevant for the evolution of dormancy and reactivation in other microbes. For instance, bacterial persistence has also been modeled as a switch between alternative states in *Escherichia coli* and different types of persister cells may be triggered by different cues to cope with transient exposition to antibiotics [3]. Dormancy and reactivation have also been described in several temperate viruses infecting bacteria [4, 5, 9, 33]. Indeed, temperate bacteriophages also need to chose between a lysogenic and a lytic state which are equivalent to the dormant and the infectious states in the present model. High density of susceptible bacteria selects for the lytic transmission routes. In contrast, when opportunities for horizontal transmissions are reduced, the lysogenic transmission route is favored. Interestingly, some viruses can use different cues to track various components of their environments and modify their phenotypes [4, 5, 13]. The fact that the genetics of the regulation of the phenotypic switch between alternative states is often well characterized in viruses [11, 14, 30] could facilitate future attempts to test the validity of our theoretical predictions on the evolution of plasticity in fluctuating environments.

Finally, it is important to note that our work focuses on periodic and deterministic fluctuations of the environment. This analysis is particularly relevant for seasonal variations that are reasonably predictable from one year to the next, but some environmental perturbations are much less predictable. Previous analysis of the evolution of dormancy have often focused on the effects of stochastic perturbations of the environment [12, 20] and it would be interesting to explore the effect of environmental stochasticity on the robustness of our predictions. Interestingly, using an indirect cue like mosquito bites, may provide a more accurate information on transmission opportunities than fixed dates based on photoperiod. We hope the present work may provide a suitable theoretical framework to explore the evolution of the plasticity in pathogen dormancy in a broader range of environmental scenarios.

## Acknowledgements

We thank Ana Rivero and Arthur Talman for many inspiring discussions and for useful comments on an earlier draft. SG ackowledges financial support from ANR grant HAMLET (ANR-23-CE02-0013).

