## Supplementary Online Material for "Evolution of pathogen dormancy in fluctuating environments"

### Supplementary Information

### S1 Epidemiological dynamics

#### S1.1 Without seasonality

We compute the reproductive ratio $R_0$ using the next generation approach after linearising the system at the disease free equilibrium $I_V = I_H = D_H = R_H = 0, N_V = S_V$ with $\mathbf{X}(t) = (I_H, D_H, I_V)^\top$ satisfies the system.

$$\frac{d\mathbf{X}(t)}{dt} = \mathbf{A}\mathbf{X}(t), \tag{S1.1}$$

where

$$\mathbf{A} = \begin{pmatrix} -(d+\gamma_I) & r & \beta_V \\ d & -(r+\gamma_D) & 0 \\ \beta_H N_V & (1-c_D)\beta_H N_V & -\mu_V \end{pmatrix}. \tag{S1.2}$$

By the representation $\mathbf{A}$ as $\mathbf{A} = \mathbf{F} - \mathbf{V}$ with

$$\mathbf{F} = \begin{bmatrix} 0 & r & \beta_V \\ d & 0 & 0 \\ \beta_H N_V & (1-c_D)\beta_H N_V & 0 \end{bmatrix}, \quad \mathbf{V} = \begin{bmatrix} (d+\gamma_I) & 0 & 0 \\ 0 & (r+\gamma_D) & 0 \\ 0 & 0 & \mu_V \end{bmatrix},$$

which satisfy the next-generation theorem [2] then the basic reproductive value $R_0$ is given by

$$R_0 = \rho\left(\mathbf{F}\mathbf{V}^{-1}\right) \text{ where } \mathbf{F}\mathbf{V}^{-1} = \begin{bmatrix} 0 & \dfrac{r}{r+\gamma_D} & \dfrac{\beta_V}{\mu_V} \\ \dfrac{d}{d+\gamma_I} & 0 & 0 \\ \dfrac{\beta_H N_V}{d+\gamma_I} & \dfrac{(1-c_D)}{\beta}{}_H N_V r+\gamma_D & 0 \end{bmatrix} \tag{S1.3}$$

Unfortunately, we do not have an analytic formula for $R_0$. But, since we know that $R_0$ is the only positive root of the characteristic polynomial $P(x) = \det(x\mathbf{I} - \mathbf{F}\mathbf{V}^{-1})$, we infer that

$$\begin{aligned} R_0 > 1 &\iff P(1) < 0 \\ &\iff \frac{\beta_V \beta_H N_V \left(r + \gamma_D + d(1-c_D)\right)}{\mu_V \left(\gamma_I + d\right)\left(r+\gamma_D\right)} + \frac{rd}{\left(\gamma_I + d\right)\left(r+\gamma_D\right)} > 1 \end{aligned}$$

#### S1.2 With Seasonality

In this section, we assume that the dormancy and recovery rate are fixed meanwhile the growth rate of vector fluctuate seasonally. We can assume that in one period the growth rate has the following step form:

$$\rho_V(t) = \begin{cases} \rho & \text{if } 0 \le \left\{\dfrac{t}{T}\right\} < 1-\sigma \\ 0 & \text{otherwise} \end{cases} \tag{S1.4}$$

##### S1.2.1 The periodic attractor of the vector population $\hat{N}_V(t)$

**Proposition S1.1.** *We consider the following ODE:*

$$\begin{cases} \dfrac{dx(t)}{dt} & = \rho_V(t)x(t)\left(1 - \dfrac{x(t)}{K}\right) - \mu_V(t)x(t) \\ x(t_0T) & = x_0 \ . \end{cases} \tag{S1.5}$$

*Where $\rho_V(t), \mu_V(t)$ are positive $T$-periodic functions. Assume that $\varphi(T) > 0$ where*

$$\varphi(t) = \int_0^t (\rho_V(s) - \mu_V(s))\, ds \, .$$

*Then, there is a unique periodic attractor given by:*

$$x^\star(t) = \frac{e^{\varphi(t)}}{\int_0^t \frac{\rho_V(s)}{K} e^{\varphi(s)} ds + \frac{1}{e^{\varphi(T)}-1} \int_0^T \frac{\rho_V(s)}{K} e^{\varphi(s)} ds}. \tag{S1.6}$$

*Proof.* If we let $y(t) = \frac{1}{x(t)}$, we see that $y$ is the solution of a linear ODE

$$\frac{dy}{dt} = -(\rho_V(t) - \mu_V(t))y(t) + \frac{\rho_V(t)}{K} \, .$$

Therefore the general solution of (S1.5) is

$$x(t) = \frac{e^{\varphi(t)}}{C + \int_0^t \frac{\rho_V(s)}{K} e^{\varphi(s)} ds} \, ,$$

with $C$ a suitable constant. Thanks to the periodicity of functions $\rho_V$ and $\mu_V$, we find that for another constant $C'$:

$$x(t + nT) = \frac{e^{\varphi(t)}}{C' e^{-n\varphi(T)} + \frac{1-e^{-n\varphi(T)}}{e^{\varphi(T)}-1} \int_0^T \frac{\rho_V(s)}{K} e^{\varphi(s)} ds + \int_0^t \frac{\rho_V(s)}{K} e^{\varphi(s)} ds} \, .$$

Therefore we obtain that $x^*$ is the unique periodic attractor, that is

$$\lim_{n \to +\infty} x(t + nT) - x^*(t) = 0 \, .$$

□

##### S1.2.2 Influence of seasonality on the growth rate $\lambda$

We linearise the system near the disease free equilibrium where $S_H = 1$, $S_V = N_V$ and $D_H = R_H = I_H = I_V = 0$.

$$\frac{d\mathbf{X}}{dt}(t) = \mathbf{A}(t)\mathbf{X} \tag{S1.7}$$

where,

$$\mathbf{A}(t) = \begin{pmatrix} -(\gamma_I + d) & r & \beta_V \\ d & -(r + \gamma_D) & 0 \\ \beta_H \hat{N}_V(t) & (1 - c_D)\beta_H \hat{N}_V(t) & -\mu_V \end{pmatrix}. \tag{S1.8}$$

**Assumption:** By the transformation $\mathbf{A}_0 \to \mathbf{A}_0 + \kappa \mathbf{I}$ translate $\lambda$ - the spectral radius of matrix $\exp(T\mathbf{A}_0)$ to $\lambda \to \lambda \exp(\kappa T)$. Without loss in generality and to simplify statements, we assume that $R_0 = \lambda_0 = 1$.

**Proposition S1.2.** *For* $\lambda = \rho\left(\Phi_{\boldsymbol{A}}(T)\right)$ *the spectral radius of the monodromy matrix we have*

$$\lambda = 1 + \sigma\lambda_1 + o(\sigma) \tag{S1.9}$$

*with* $\lambda_1 < 0$ *therefore seasonality decreases the growth rate of the population.*

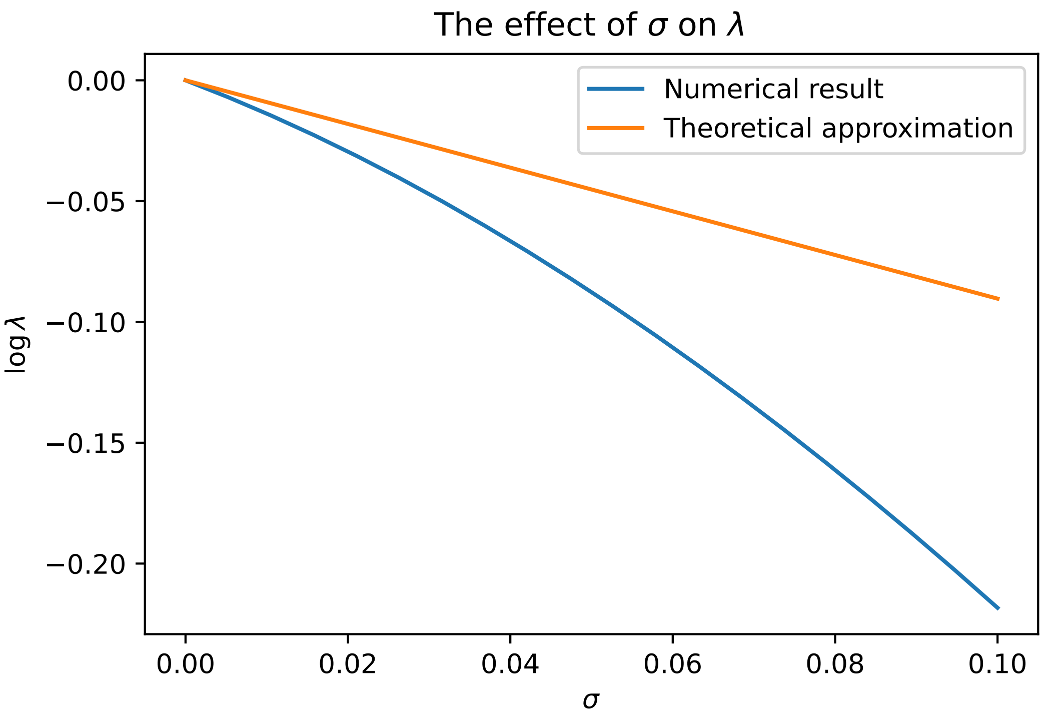


**Figure S1: The effect of seasonality on the persistence of the pathogen.** An increase of $\sigma$ (the duration of the low-transmission season) has a negative effect on pathogen persistence measured by the growth rate $\lambda$. Parameters: $T = 5, \alpha = 0, \sigma = 0.5, \beta_V = 1, \beta_H = 1, \gamma = 2/3, \mu_V = 1, \mu_H = 0.5, d = 1, r = 1, \rho = 4, K = 1$.

*Proof.* From the explicit formula (2.2) we infer the Taylor expansion

$$\begin{aligned}\hat{N}_V(t) &= \frac{K(\rho-\mu_V)}{\rho}\left(1 - \frac{\sigma T \mu_V e^{T(\rho-\mu_V)}}{e^{T(\rho-\mu_V)}-1}\right) + o(\sigma) && \text{if } 0 < t < (1-\sigma)T \\ &= \frac{K(\rho-\mu_V)}{\rho}\left(1 - \frac{\sigma T \mu_V e^{T(\rho-\mu_V)}}{e^{T(\rho-\mu_V)}-1} e^{-t(\rho-\mu_V)}\right) + o(\sigma) && \text{if } (1-\sigma)T < t < 1\end{aligned}$$

We can rewrite $\mathbf{A}(t)$ as the following form. $\mathbf{A}(t) = \mathbf{A_0} + \mathbf{A_1} e^{-T\left\{\frac{t}{T}\right\}(\rho-\mu_V)}\sigma + o(\sigma)$ where $\alpha = \beta_H K \frac{\rho-\mu_V}{\rho}$

$$\mathbf{A_0} = \begin{pmatrix} -(\gamma_I + d) & r & \beta_V \\ d & -(r+\gamma_D) & 0 \\ \alpha & (1-c_D)\alpha & -\mu_V \end{pmatrix} \tag{S1.10}$$

and

$$\mathbf{A_1} = -T\mu_V\beta_H\alpha\frac{e^{T(\rho-\mu_V)}}{e^{T(\rho-\mu_V)}-1}\begin{pmatrix} 0 & 0 & 0 \\ 0 & 0 & 0 \\ 1 & 1-c_D & 0 \end{pmatrix} \tag{S1.11}$$

Let $\mathbf{L} = \Phi_{\mathbf{A}}(T)$ be the monodromy matrix associated to matrix $A(t)$. We assume that $v_0, u_0$ are the left(right)eigenvector associated to matrix $A_0$. We use Duhamel's formula with $\mathbf{L} = \mathbf{L_0} + \sigma\mathbf{L_1} + o(\sigma)$ where

$$\mathbf{L_0} = e^{T\mathbf{A_0}} \tag{S1.12}$$

$$\mathbf{L_1} = \int_0^T v_0 e^{(T-s)\mathbf{A_0}}\mathbf{A_1}e^{s\mathbf{A_0}}u_0 e^{-s(\rho-\mu_V)}ds \tag{S1.13}$$

Combining this Taylor expansion with the expansion $\lambda = 1+\sigma\lambda_1+o(\sigma)$ and the equation $\mathbf{L}u = \lambda u$ we obtain, since $v_0e^{tA_0} = v_0$ and $e^{tA_0}u_0 = u_0$,

$$\lambda_1 = v_0\mathbf{L_1}u_0 = \int_0^T v_0 e^{(T-s)\mathbf{A_0}}\mathbf{A_1}e^{s\mathbf{A_0}}u_0 e^{-s(\rho-\mu_V)}\,ds = v_0\mathbf{A_1}u_0\int_0^T e^{-s(\rho-\mu_V)}\,ds. \tag{S1.14}$$

Since the integral factor is positive, since $u_0$ and $v_0$ have positive entries, and since $\mathbf{A_1}$ has zero or negative entries, we obtain that $\lambda_1 < 0$. Tedious but straightforward computations of $u_0, v_0$ gives the exact formula which we used in **Figure S1**:

$$\lambda_1 = -T\mu_V\alpha\frac{\beta_V}{\mu_V}\left(1+(1-c_D)\frac{d}{r+\gamma_D}\right)\left(1+\kappa\frac{d}{r+\gamma_D}+\delta\frac{\beta_V}{\mu_V}\right)^{-1} \tag{S1.15}$$

with

$$\delta = \frac{(d+\gamma_I)(r+\gamma_D)-rd}{r+\gamma_D}, \quad \kappa = \left(\frac{r}{r+\gamma_D}+(1-c_D)\alpha\frac{\beta_V}{\mu_V}\right) \tag{S1.16}$$

□

The above derivation indicates that seasonality $\sigma$ decreases the persistence of the pathogen. This is due to the effect of seasonality on the reduction of the average density of the mosquito vector. Indeed we can show that:

**Proposition S1.3.** *The mean of vector population is given by:*

$$\left\langle \hat{N}_V \right\rangle = \frac{1}{T}\left(\frac{K(\rho-\mu)}{\rho}(1-\sigma)T - \frac{K\sigma T\mu}{\rho} + \frac{K(\rho-\mu)}{\rho}\frac{\tau_1-1}{\tau_2-1}\frac{\left(e^{\sigma T\mu}-1\right)}{\mu}\right) \tag{S1.17}$$

*with* $\tau_1 = e^{T((1-\sigma)\rho-\mu)}$ *and* $\tau_2 = e^{T(1-\sigma)(\rho-\mu)}$ *and it decreases with increasing seasonality.*

*Proof.* The formula is derived by integrating the formula (2.2). Then we differentiate the expression with respect to $\sigma$ to obtain the monotonicity. □

### S2 Evolutionary dynamics

As discussed in the main text, the selection gradient on a trait $z$ is given by equation (3.2) (which depends on the dynamics of the frequency $f_k(t)$ and the reproductive value

$v_j(t)$ which are given below.

**Class frequencies:**
The dynamics of class frequencies is given by:

$$\frac{df_{I_H}(t)}{dt} = -\left(d+\gamma_I\right) f_{I_H}(t) + r f_{D_H}(t) + \beta_V \hat{S}_H(t) f_{I_V}(t) - r_w f_{I_H}(t) \tag{S2.1}$$
$$\frac{df_{D_H}(t)}{dt} = d\ f_{I_H}(t) - \left(r+\gamma_D\right) f_{D_H}(t) - r_w f_{D_H}(t) \tag{S2.2}$$
$$\frac{df_{I_V}(t)}{dt} = \beta_H \hat{S}_V(t) f_{I_H}(t) + (1-c_D)\beta_H \hat{S}_V(t) f_{D_H}(t) - \mu_V f_{I_V}(t) - r_w f_{I_V}(t) \tag{S2.3}$$

Where $r_w = \sum_{j,k} a_w^{jk}(t) f^k(t)$ is the per-capital growth rate of resident population. When the system sits on its periodic attractor $\langle r_w \rangle = 0$ and $\left\langle \frac{d}{dt} \ln\left(f_k\right) \right\rangle = 0$. If we divide both side of (S2.2) by $f_{D_H}$ and average over one period we can show that:

$$\left\langle \frac{\hat{f}_{I_H}}{\hat{f}_{D_H}} \right\rangle = \frac{r+\gamma_D}{d}. \tag{S2.4}$$

**Reproductive values:**
The dynamics of reproductive values are given by:

$$\frac{dv_{I_H}(t)}{dt} = \left(\gamma_I + d\right) v_{I_H}(t) - d v_{D_H}(t) - \beta_H \hat{S}_V(t) v_{I_V}(t) + r_w v_{I_H}(t) \tag{S2.5}$$
$$\frac{dv_{D_H}(t)}{dt} = -r v_{I_H}(t) + \left(r+\gamma_D\right) v_{D_H}(t) - (1-c_D)\beta_H \hat{S}_V(t) v_{I_V}(t) + r_w v_{D_H}(t) \tag{S2.6}$$
$$\frac{dv_{I_V}(t)}{dt} = -\beta_V \hat{S}_H(t) v_{I_H}(t) + \mu_V v_{I_V}(t) + r_w v_{I_V}(t) \tag{S2.7}$$

When the system sits on its periodic attractor $\langle r_w \rangle = 0$ and $\left\langle \frac{d}{dt} \ln\left(v_k\right) \right\rangle = 0$. If we divide both side of (S2.7) by $v_{I_V}$ and average over one period we can show that

$$\left\langle \frac{\hat{S}_H \hat{v}_{I_H}}{\hat{v}_{I_V}} \right\rangle = \frac{\mu_V}{\beta_V}. \tag{S2.8}$$

#### S2.1 Evolution without seasonality

In the absence of seasonality, the condition (S2.8) yields: $\hat{v}_{I_V} = \hat{v}_{I_H} \beta_V \hat{S}_H / \mu_V$ and we can then use (S2.6) to show that:

$$\hat{S}_V = \frac{\left(r(\hat{v}_{D_H} - \hat{v}_{I_H}) + \hat{v}_{D_H}\gamma_D\right)\mu_V}{\hat{S}_H v_{I_H}(1-c_D)\beta_H\beta_V}. \tag{S2.9}$$

Using this expression of $\hat{S}_V$ in (S2.5) yields:

$$\hat{v}_{I_H} = \hat{v}_{D_H} \frac{(r+\gamma_D) + d(1-c_D)}{r + (d+\gamma_I)(1-c_D)}. \tag{S2.10}$$

The selection gradient on dormancy (3.3) shows that dormancy will be favored when $\hat{v}_{D_H}/\hat{v}_{I_H} > 1$. Hence, dormancy is favored when $c_D < c_D^* = 1 - \frac{\gamma_D}{\gamma_I}$. Interestingly, the selection gradient on reactivation (3.4) yields the opposite prediction: reactivation is selected for when dormancy is not. In other words, in the absence of seasonality, dormancy evolves when $c_D < c_D^*$, and reactivation evolves when $c_D > c_D^*$.

#### S2.2 Evolution with seasonality

In the absence of plasticity we use expressions (3.3) and (3.4) to plot the gradient vector field $(\langle \mathcal{S}_d \rangle, \langle \mathcal{S}_r \rangle)$ for a given cost of dormancy. The coevolutionary stable strategy $(d^*, r^*)$ is the point where both selection gradient vanish. Numerical simulations indicate that this coESS is unique over the range of parameter values we considered (see **Figure S2**).

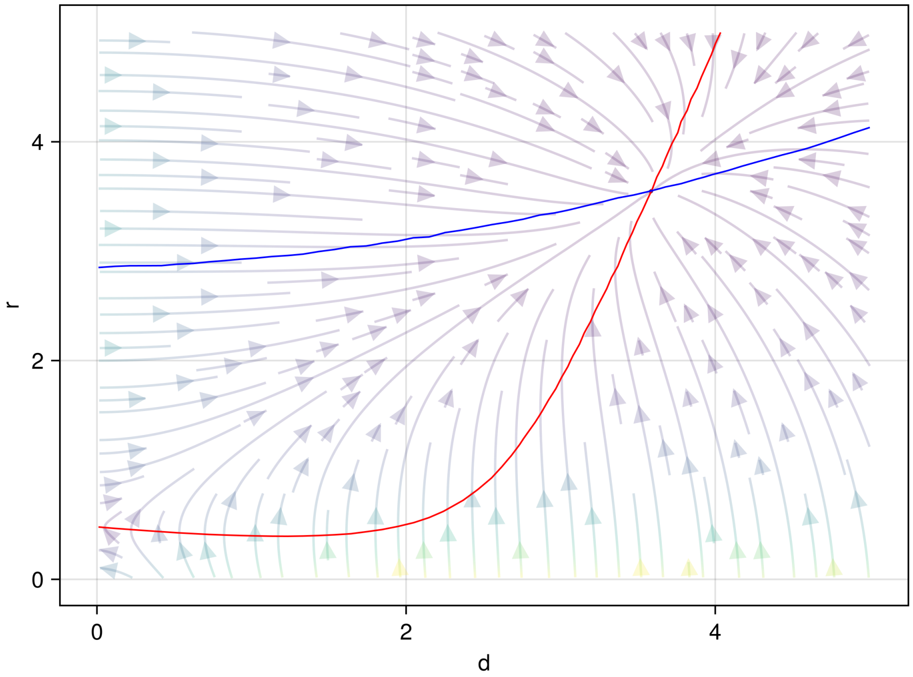


**Figure S2: Finding the coESS for $d$, $r$.** The coESS coincides with the intersection of the null-clines (i.e., where both the selection gradients on $d$ and $r$ vanish): the red curve is $\langle \mathcal{S}_d \rangle = 0$, the blue curve is $\langle \mathcal{S}_r \rangle = 0$. Parameters : $c_D = 0.92$.

In contrast to the scenario without seasonality, we find that the coESS can lead to intermediate values $d^*$ and $r^*$ (see **Figures S3-S6**). To understand the influence of seasonality on the coESS the selection coefficients can be rewritten as:

$$\langle \mathcal{S}_d \rangle > 0 \implies \langle \hat{v}_{I_H} - \hat{v}_{D_H} \rangle < -\frac{\mathrm{cov}\left(\hat{f}_{I_H}, (\hat{v}_{I_H} - \hat{v}_{D_H})\right)}{\left\langle \hat{f}_{I_H} \right\rangle} \tag{S2.11}$$

$$\langle \mathcal{S}_r \rangle > 0 \implies \langle \hat{v}_{I_H} - \hat{v}_{D_H} \rangle > -\frac{\mathrm{cov}\left(\hat{f}_{D_H}, (\hat{v}_{I_H} - \hat{v}_{D_H})\right)}{\left\langle \hat{f}_{D_H} \right\rangle} \tag{S2.12}$$

Hence, in a seasonal environments, the selection for $d$ and for $r$ ar not necessarily of opposite signs because of the covariance between the epidemiological dynamics (i.e. $\hat{f}_{I_H}$ or $\hat{f}_{D_H}$) and the fluctuations of $\hat{v}_{I_H} - \hat{v}_{D_H}$.

Next, we consider a scenario where the pathogen is allowed to adopt a plastic strategy where its phenotypes can change during the year. We consider two scenarios depending on the cue used by the pathogen to change its phenotype: (1) the cue is time (i.e. the switch of the phenotype is triggered at specific dates during the year), (2) the cue is vector vector density (i.e. the switch of the phenotype is triggered when the vector density reaches a specific threshold).

##### S2.2.1 Plasticity with time

**Selection for plastic dormancy:**
Here we assume that dormancy $d(t)$ fluctutates with time

$$d(t) = \begin{cases} d_{\text{on}} & \text{if } t \in [t_s^d, t_e^d] \\ d_{\text{off}} = 0 & \text{otherwise} \end{cases} \tag{S2.13}$$

We study the evolution of the phenotypes $d_{\text{on}}$, $t_s^d$ and $t_e^d$ using the chain rule in the definition of the instantaneous gradient (3.2)

$$\frac{\partial \mathbf{A}_m}{\partial d_{\text{on}}} = \frac{\partial \mathbf{A}_m}{\partial d}\frac{\partial d(t)}{\partial d_{\text{on}}} = \begin{pmatrix} -1 & 0 & 0 \\ 1 & 0 & 0 \\ 0 & 0 & 0 \end{pmatrix} \mathbb{1}_{[t_s^d, t_e^d)}(t)\,. \tag{S2.14}$$

Therefore the selection gradient on $d_{\text{on}}$ is

$$\mathcal{S}_{d_{\text{on}}} = -\frac{1}{T}\int_0^T \left(\hat{v}_{I_H} - \hat{v}_{D_H}\right) \hat{f}_{I_H}(t) \mathbb{1}_{[t_s^d, t_e^d)} dt = -\frac{1}{T}\int_{t_s^d}^{t_e^d} \left(\hat{v}_{I_H} - \hat{v}_{D_H}\right) \hat{f}_{I_H}(t) dt\,. \tag{S2.15}$$

When we compute gradients with respect to $t_s^d$ and $t_e^d$, we have to introduce the Dirac distribution $\delta_a$ such that $\int f(t)\delta_a(dt) = f(a)$. Indeed, the derivative, in the sense of distributions, of the Heaviside function $H(t) = \mathbf{1}_{(t\geq 0)}$ is $\delta_0 = \dfrac{\partial H(t)}{\partial t}$ and $\delta_a = -\dfrac{\partial H(t-a)}{\partial a}$. Hence, we can use this Heaviside function to rewrite (S2.13) in the following way

$$d(t) = d_{\text{on}} \left(H(t - t_s^d) - H(t - t_e^d)\right)$$

we obtain that the partial derivatives, in the sense of distributions, of $d(t)$ are

$$\frac{\partial d(t)}{\partial t_s^d} = -d_{\text{on}} \delta_{t_s^d}(dt) \tag{S2.16}$$

$$\frac{\partial d(t)}{\partial t_e^d} = d_{\text{on}} \delta_{t_e^d}(dt). \tag{S2.17}$$

Alternatively, one may approximate the function $H(x)$ with the logistic function $H_k(x) = (1 + e^{-2kx})^{-1}$ with a large value of $k$. This may allow us to consider more continuous and perhaps more realistic transitions of the phenotypes (the larger the value of $k$, the sharper the phenotypic switch). This alternative formalism, however, yields a slightly more complex selection gradient which depends on $k$ (see [1]). Hence, in the present work we consider the limit case where $k \to \infty$ which is equivalent to a discontinuous switch and, using (S2.16) and (S2.17), yields the following selection gradient on $t_s^d$ and $t_e^d$, respectively:

$$\mathcal{S}_{t_s^d} = \frac{1}{T}\int_0^T d_{\text{on}}\left(\hat{v}_{I_H} - \hat{v}_{D_H}\right)\hat{f}_{I_H}(t)\delta_{t_s^d}(dt) \tag{S2.18}$$

$$= \frac{d_{\text{on}}\hat{f}_{I_H}(t_s^d)}{T}\left(\hat{v}_{I_H}(t_s^d) - \hat{v}_{D_H}(t_s^d)\right) \tag{S2.19}$$

$$\mathcal{S}_{t_e^d} = -\frac{1}{T}\int_0^T d_{\text{on}}\left(\hat{v}_{I_H} - \hat{v}_{D_H}\right)\hat{f}_{I_H}(t)\delta_{t_e^d}(dt) \tag{S2.20}$$

$$= -\frac{d_{\text{on}}\hat{f}_{I_H}(t_e^d)}{T}\left(\hat{v}_{I_H}(t_e^d) - \hat{v}_{D_H}(t_e^d)\right) \tag{S2.21}$$

**Selection for plastic reactivation:**
Here we assume that reactivation $r(t)$ fluctutates with time

$$r(t) = \begin{cases} r_{\text{on}} & \text{if } t \in [t_s^r, t_e^r] \\ r_{\text{off}} = 0 & \text{otherwise} \end{cases} \tag{S2.22}$$

Following the same approach as above, we study the evolution of the phenotypes $r_{\text{on}}$, $t_s^r$ and $t_e^r$ and we obtain:

$$\mathcal{S}_{r_{\text{on}}} = \frac{1}{T}\int_0^T \left(\hat{v}_{I_H} - \hat{v}_{D_H}\right)\hat{f}_{D_H}(t)\mathbb{1}_{[t_s^r, t_e^r)}dt \tag{S2.23}$$

$$= \frac{1}{T}\int_{t_s^r}^{t_e^r} \left(\hat{v}_{I_H} - \hat{v}_{D_H}\right)\hat{f}_{D_H}(t)dt \tag{S2.24}$$

$$\mathcal{S}_{t_s^r} = -\frac{1}{T}\int_0^T r_{\text{on}}\left(\hat{v}_{I_H} - \hat{v}_{D_H}\right)\hat{f}_{D_H}(t)\delta_{t_s^r}(dt) \tag{S2.25}$$

$$= -\frac{r_{\text{on}}\hat{f}_{D_H}(t_s^r)}{T}\left(\hat{v}_{I_H}(t_s^r) - \hat{v}_{D_H}(t_s^r)\right) \tag{S2.26}$$

$$\mathcal{S}_{t_e^r} = \frac{1}{T}\int_0^T r_{\text{on}}\left(\hat{v}_{I_H} - \hat{v}_{D_H}\right)\hat{f}_{D_H}(t)\delta_{t_e^r}(dt) \tag{S2.27}$$

$$= \frac{r_{\text{on}}\hat{f}_{D_H}(t_e^r)}{T}\left(\hat{v}_{I_H}(t_e^r) - \hat{v}_{D_H}(t_e^r)\right) \tag{S2.28}$$

In the following, we assume for simplicity that the phenotypes $d_{\text{on}}$ and $r_{\text{on}}$ are fixed and we focus on the evolution of the timing of the switch of dormancy and reactivation. Let us assume that there is an ESS with respect to $t_s^d, t_e^d, t_s^r$ and $t_e^r$ so that $\mathcal{S}_{t_s^d} = \mathcal{S}_{t_e^d} = \mathcal{S}_{t_s^r} = \mathcal{S}_{t_e^r} = 0$. The above selection gradients show that the ESS condition implies that switching times take place when $\hat{v}_{I_H}(t) = \hat{v}_{D_H}(t)$ such that $\hat{v}_{I_H}(t) - \hat{v}_{D_H}(t)$ is positive before $t_s^d$ and negative after $t_s^d$ (is negative before $t_e^d$ and positive after $t_e^d$). Hence the switching times for dormancy and reactivation are the same but dormancy is switched "on" when reactivation is switched "off", and vice versa. Hence, the evolution of the pathogen is expected to yield a "bang-bang" strategy characterized by a switch to a dormant state when it encounters an unfavorable environment, until the next favorable period where it reactivates (see **Figure 4**).

##### S2.2.2 Plasticity with vector density

Here we assume that dormancy $d(t)$ and reactivation $r(t)$ fluctuate with time

$$d(t) = \begin{cases} d_{\text{on}} & \text{if } N_V(t) < N_V^* \\ d_{\text{off}} = 0 & \text{otherwise} \end{cases} \tag{S2.29}$$

$$r(t) = \begin{cases} r_{\text{on}} & \text{if } N_V(t) > N_V^* \\ r_{\text{off}} = 0 & \text{otherwise} \end{cases} \tag{S2.30}$$

Under this scenario the two quantities $t_s^d = t_e^r, t_e^d = t_s^r$ depend on the threshold value $N_V^*$. We study below the selection acting on this threshold value.

$$\mathcal{S} = \left\langle v \frac{\partial \mathbf{A}}{\partial N_V^*}(t) f \right\rangle, \text{ with } \frac{\partial \mathbf{A}}{\partial N_V^*}(t) = \frac{\partial \mathbf{A}}{\partial d}(t) \frac{\partial d}{\partial N_V^*}(t) + \frac{\partial \mathbf{A}}{\partial r}(t) \frac{\partial r}{\partial N_V^*}(t) \tag{S2.31}$$

we have

$$\frac{\partial d}{\partial N_V^*}(t) = -d_{\text{on}} \frac{\partial t_s^d}{\partial N_V^*}(t) \delta_{t_s^d}(dt) + d_{\text{on}} \frac{\partial t_e^d}{\partial N_V^*}(t) \delta_{t_e^d}(dt), \tag{S2.32}$$

$$\frac{\partial r}{\partial N_V^*}(t) = r_{\text{on}} \frac{\partial t_s^d}{\partial N_V^*}(t) \delta_{t_s^d}(dt) - r_{\text{on}} \frac{\partial t_e^d}{\partial N_V^*}(t) \delta_{t_e^d}(dt) \tag{S2.33}$$

We compute $\frac{\partial t_s^d}{\partial N_V^*}(t)$ from the equation $\hat{N}_V(t_s^d) = N_V^*$ that we differentiate

$$1 = \frac{\partial N_V^*}{\partial N_V^*} = N_V^{'}(t_s^d) \frac{\partial t_s^d}{\partial N_V^*}(t)$$

and since $N_V^{'}(t) = \left( \rho_V(t) \left( 1 - \frac{\hat{N}_V(t)}{K} \right) - \mu_V \right) \hat{N}_V(t)$ we get

$$\frac{\partial t_s^d}{\partial N_V^*}(t) = \frac{1}{N_V^{'}(t_s^d)} = -\frac{1}{\mu_V N_V^*} < 0, \quad \frac{\partial t_e^d}{\partial N_V^*}(t) = \frac{1}{N_V^{'}(t_e^d)} = \frac{1}{\left( \rho \left( 1 - \frac{N_V^*}{K} \right) - \mu_V \right) N_V^*} > 0. \tag{S2.34}$$

Note that the signs agree with the fact that at $t = t_s^d$ the curve $N_V(t)$ is crossing downward the level $N_V^*$, while at $t = t_e^d$ the curve $N_V(t)$ is crossing upward the level $N_V^*$. Putting everything together gives the gradient $\mathcal{S} = \mathcal{S}_{N_V^*} = \langle \mathcal{S}(t) \rangle$ with

$$\mathcal{S}(t) = -\left( \hat{v}_{I_H} - \hat{v}_{D_H} \right) \left( \frac{\partial d}{\partial N_V^*}(t) \hat{f}_{I_H} - \frac{\partial r}{\partial N_V^*}(t) \hat{f}_{D_H} \right) \tag{S2.35}$$

$$= -\left( \hat{v}_{I_H} - \hat{v}_{D_H} \right) \times \tag{S2.36}$$

$$\left( \hat{f}_{I_H} d_{\text{on}} \left( \frac{\partial t_e^d}{\partial N_V^*}(t) \delta_{t_e^d}(dt) - \frac{\partial t_s^d}{\partial N_V^*}(t) \delta_{t_s^d}(dt) \right) - \hat{f}_{D_H} r_{\text{on}} \left( \frac{\partial t_s^d}{\partial N_V^*}(t) \delta_{t_s^d}(dt) - \frac{\partial t_e^d}{\partial N_V^*}(t) \delta_{t_e^d}(dt) \right) \right) \tag{S2.37}$$

$$= -\left( \hat{v}_{I_H} - \hat{v}_{D_H} \right) \times \tag{S2.38}$$

$$\left(-\left(\hat{f}_{I_H} d_{\text{on}} + \hat{f}_{D_H} r_{\text{on}}\right) \frac{\partial t_s^d}{\partial N_V^*}(t)\delta_{t_s^d}(dt) + \left(\hat{f}_{I_H} d_{\text{on}} + \hat{f}_{D_H} r_{\text{on}}\right) \frac{\partial t_e^d}{\partial N_V^*}(t)\delta_{t_e^d}(dt)\right) \tag{S2.39}$$

Therefore, after integrating the gradient of selection over one period we obtain:

$$\begin{aligned} \mathcal{S} = &- \left(\hat{v}_{I_H}(t_s^d) - \hat{v}_{D_H}(t_s^d)\right) \left(\hat{f}_{I_H}(t_s^d) d_{\text{on}} + \hat{f}_{D_H}(t_s^d) r_{\text{on}}\right) \frac{1}{\mu_V N_V^*} \\ &- \left(\hat{v}_{I_H}(t_e^d) - \hat{v}_{D_H}(t_e^d)\right) \left(\hat{f}_{I_H}(t_e^d) d_{\text{on}} + \hat{f}_{D_H}(t_e^d) r_{\text{on}}\right) \frac{1}{\left(\rho\left(1 - \frac{N_V^*}{K}\right) - \mu_V\right) N_V^*} \end{aligned} \tag{S2.40}$$

We compute numerically the gradient $\mathcal{S}$ to obtain the evolutionary stable threshold $N_V^*$ using the following procedure. Given a threshold value $N_V^*$, $t_s^d$ is the time where the curve $t \to N_V(t)$ crosses the level $N_V^*$ downwards, and $t_e^d$ is the time where it crosses it upwards. Then we plug these values into (S2.40). By a numerical procedure we find the optimal $N_V^*$ such that $\mathcal{S} = 0$. The results of simulations are shown in **Figure 4**. In this scenario, pathogens enter a state of dormancy in response to an unfavorable period, guided by cues from the density of the number vector. Consequently, the dormancy period occurs later in the winter and extends until the beginning of summer. It differs from the previous scenario (section S2.2.1), where pathogens always proactively switch to dormancy before the winter.

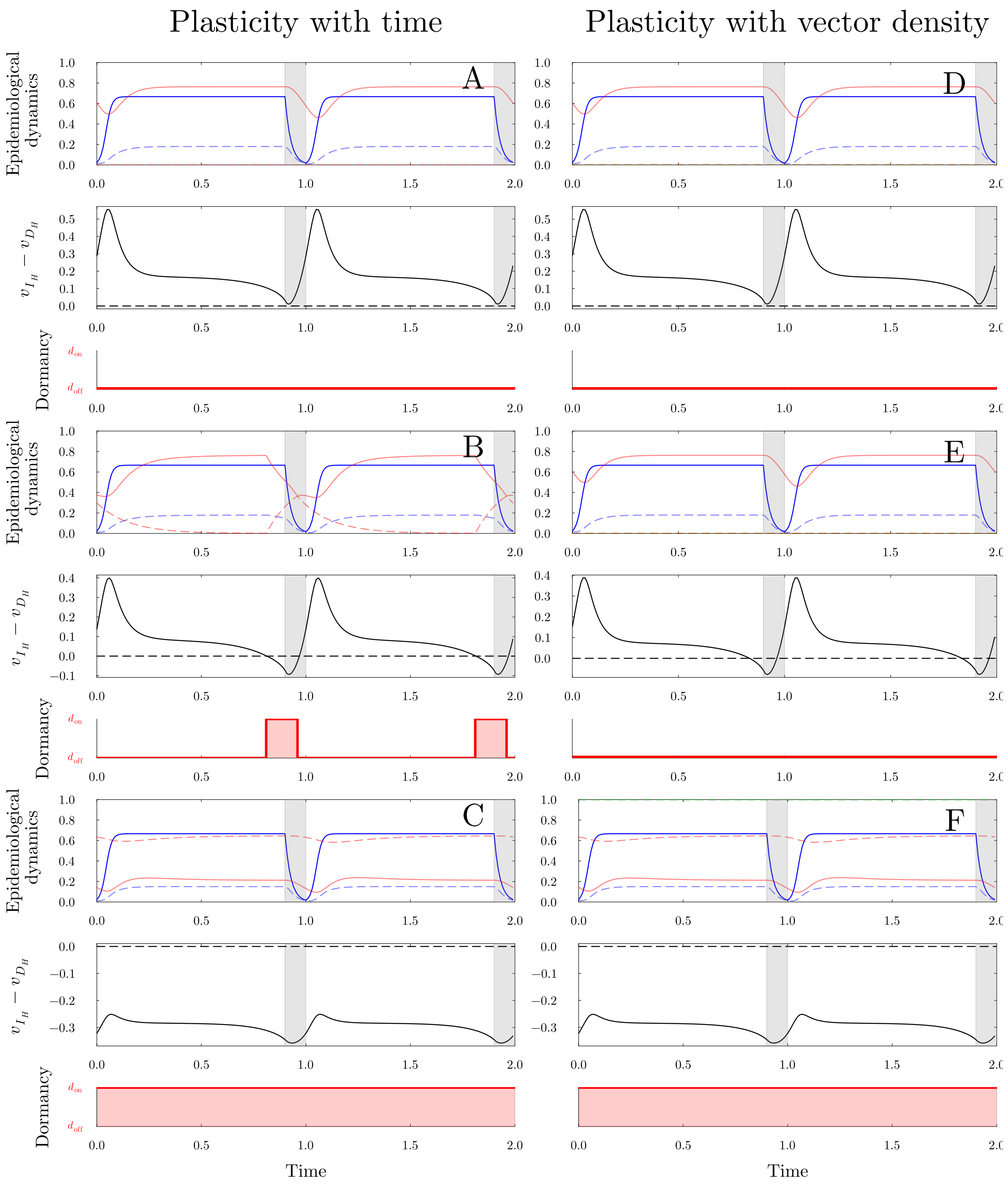


**Figure S3: Evolution of time-varying dormancy in a seasonal environment:** same as **Figure 4** but with $\sigma = 0.1$. We vary the cost of dormancy: $c_D = 1$ in (A) and (D), $c_D = 0.85$ in (B) and (E), $c_D = 0.4$ in (C) and (F). Other parameter values are given in **Table 1**.

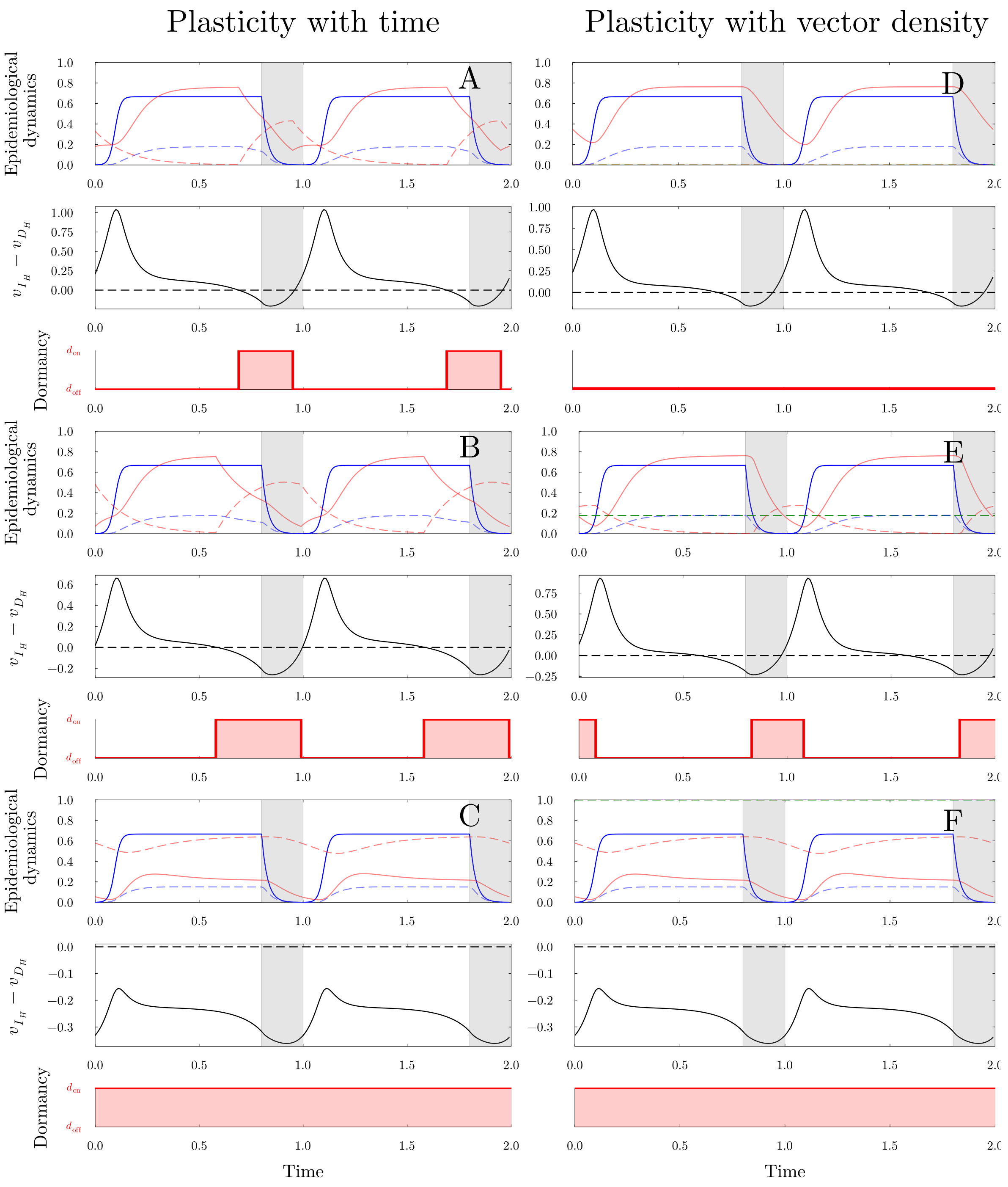


**Figure S4: Evolution of time-varying dormancy in a seasonal environment:** ame as **Figure 4** but with $\sigma = 0.2$. We vary the cost of dormancy: $c_D = 1$ in (A) and (D), $c_D = 0.85$ in (B) and (E), $c_D = 0.4$ in (C) and (F). Other parameter values are given in **Table 1**.

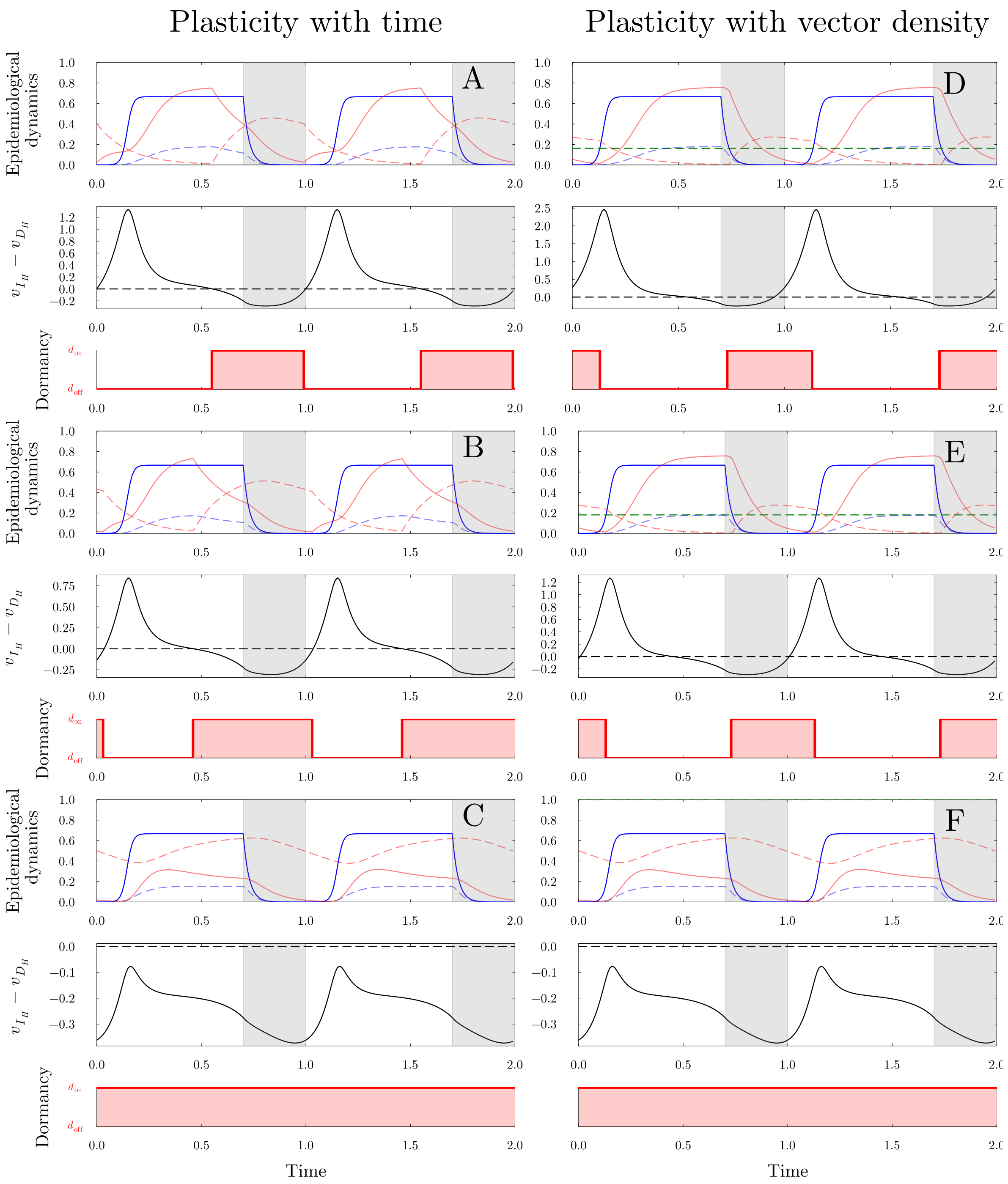


**Figure S5: Evolution of time-varying dormancy in a seasonal environment:** same as **Figure 4** but with $\sigma = 0.3$. We vary the cost of dormancy: $c_D = 1$ in (A) and (D), $c_D = 0.85$ in (B) and (E), $c_D = 0.4$ in (C) and (F). Other parameter values are given in **Table 1**.

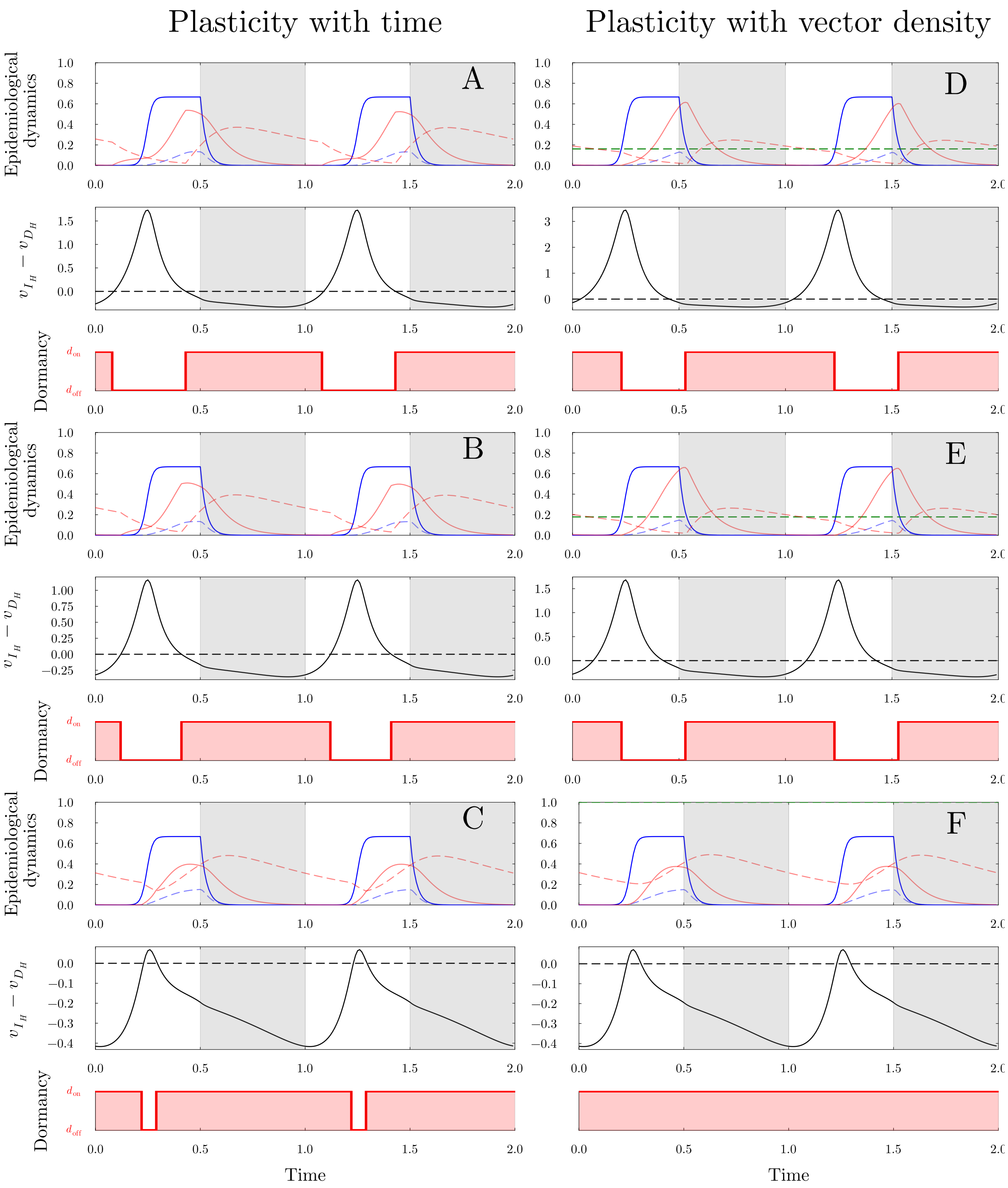


**Figure S6: Evolution of time-varying dormancy in a seasonal environment:** same as **Figure 4** but with $\sigma = 0.5$. We vary the cost of dormancy: $c_D = 1$ in (A) and (D), $c_D = 0.85$ in (B) and (E), $c_D = 0.4$ in (C) and (F). Other parameter values are given in **Table 1**.

#### S2.3 Plasticity with vector density for both dormancy and reactivation

We now assume that dormancy happens when the density of vectors $N_V(t)$ is below the level $N_V^{d*}$, and that reactivation happens when the density of vectors $N_V(t)$ is above the level $N_V^{r*}$. more precisely

$$d(t) = \begin{cases} d_{\text{on}} & \text{if } N_V(t) < N_V^{d*} \\ d_{\text{off}} = 0 & \text{otherwise} \end{cases} \tag{S2.41}$$

$$r(t) = \begin{cases} r_{\text{on}} & \text{if } N_V(t) > N_V^{r*} \\ r_{\text{off}} = 0 & \text{otherwise} \end{cases} \tag{S2.42}$$

Under this scenario $t_s^d, t_e^d$ depend on $N_V^{d*}$ and $t_s^r, t_e^r$ depend on $N_V^{r*}$. Accordingly,

$$\begin{aligned}
\mathcal{S}_{N_V^{d*}} &= \left\langle \mathbf{v}(t)\frac{\partial \mathbf{A}}{\partial N_V^{d*}}(t)\mathbf{f}(t)\right\rangle = \left\langle \mathbf{v}(t)\frac{\partial \mathbf{A}}{\partial d}(t)\mathbf{f}(t)\,\frac{\partial d}{\partial N_V^{d*}}(t)\right\rangle \\
&= \left\langle f_{I_H}(t)(v_{D_H}(t) - v_{I_H}(t))d_{\text{on}}\left(\frac{\partial t_e^d}{\partial N_V^{d*}}\delta_{t_e^d}(dt) - \frac{\partial t_s^d}{\partial N_V^{d*}}\delta_{t_s^d}(dt)\right)\right\rangle \\
&= d_{\text{on}}\left[f_{I_H}(t_e^d)(v_{D_H}(t_e^d) - v_{I_H}(t_e^d))\frac{\partial t_e^d}{\partial N_V^{d*}} - f_{I_H}(t_s^d)(v_{D_H}(t_s^d) - v_{I_H}(t_s^d))\frac{\partial t_s^d}{\partial N_V^{d*}}\right] \\
\mathcal{S}_{N_V^{r*}} &= \left\langle \mathbf{v}(t)\frac{\partial \mathbf{A}}{\partial N_V^{r*}}(t)\mathbf{f}(t)\right\rangle = \left\langle \mathbf{v}(t)\frac{\partial \mathbf{A}}{\partial r}(t)\mathbf{f}(t)\,\frac{\partial r}{\partial N_V^{r*}}(t)\right\rangle \\
&= \left\langle f_{D_H}(t)(v_{I_H}(t) - v_{D_H}(t))r_{\text{on}}\left(\frac{\partial t_e^r}{\partial N_V^{r*}}\delta_{t_e^r}(dt) - \frac{\partial t_s^r}{\partial N_V^{r*}}\delta_{t_s^r}(dt)\right)\right\rangle \\
&= r_{\text{on}}\left[f_{D_H}(t_e^r)\left(v_{I_H}(t_e^r) - v_{D_H}(t_e^r)\right)\frac{\partial t_e^r}{\partial N_V^{r*}} - f_{D_H}(t_s^r)\left(v_{I_H}(t_s^r) - v_{D_H}(t_s^r)\right)\frac{\partial t_s^r}{\partial N_V^{r*}}\right].
\end{aligned}$$

The derivatives $\frac{\partial t_s^d}{\partial N_V^{d*}}$ and $\frac{\partial t_e^d}{\partial N_V^{d*}}$ are computed as in the preceding section (see equation (S2.34)). Similarly, we obtain

$$\frac{\partial t_s^r}{\partial N_V^{r*}}(t) = \frac{1}{N_V'(t_s^r)} = \frac{1}{\left(\rho\left(1 - \frac{N_V^{r*}}{K}\right) - \mu_V\right)N_V^{r*}} > 0, \quad \frac{\partial t_e^r}{\partial N_V^{*}}(t) = \frac{1}{N_V'(t_e^r)} = -\frac{1}{\mu_V N_V^{r*}} < 0. \tag{S2.43}$$

Note that the signs are opposite of (S2.34) because at $t = t_s^r$ the curve $N_V(t)$ is crossing upward the level $N_V^{r*}$, while at $t = t_e^r$ the curve $N_V(t)$ is crossing downward the level $N_V^{r*}$. Finally, putting everything together we get the following two selection gradients:

$$\mathcal{S}_{N_V^{d*}} = -\frac{d_{\text{on}}}{N_V^{d*}}\left[f_{I_H}(t_e^d)(v_{I_H}(t_e^d) - v_{D_H}(t_e^d))\frac{1}{\rho(1 - \frac{N_V^{d*}}{K}) - \mu_V} + f_{I_H}(t_s^d)(v_{I_H}(t_s^d) - v_{D_H}(t_s^d))\frac{1}{\mu_V}\right]$$

$$\mathcal{S}_{N_V^{r*}} = -\frac{r_{\text{on}}}{N_V^{r*}}\left[f_{D_H}(t_e^r)(v_{I_H}(t_e^r) - v_{D_H}(t_e^r))\frac{1}{\mu_V} + f_{D_H}(t_s^r)(v_{I_H}(t_s^r) - v_{D_H}(t_s^r))\frac{1}{\rho(1 - \frac{N_V^{r*}}{K}) - \mu_V}\right]$$

We compute, for a given cost of dormancy $c_D = 0.8$, and duration of winter $\sigma = 0.4$, the coESS, that is the couple of values $(N_V^{d*}, N_V^{r*})$ such that $\mathcal{S}_{N_V^{d*}} = \mathcal{S}_{N_V^{r*}} = 0$. In **Figure S7** we plot the Epidemiological dynamics for the coESS, as well as the reproductive values, and finally we identify the dormancy periods in red, and the reactivation in green.

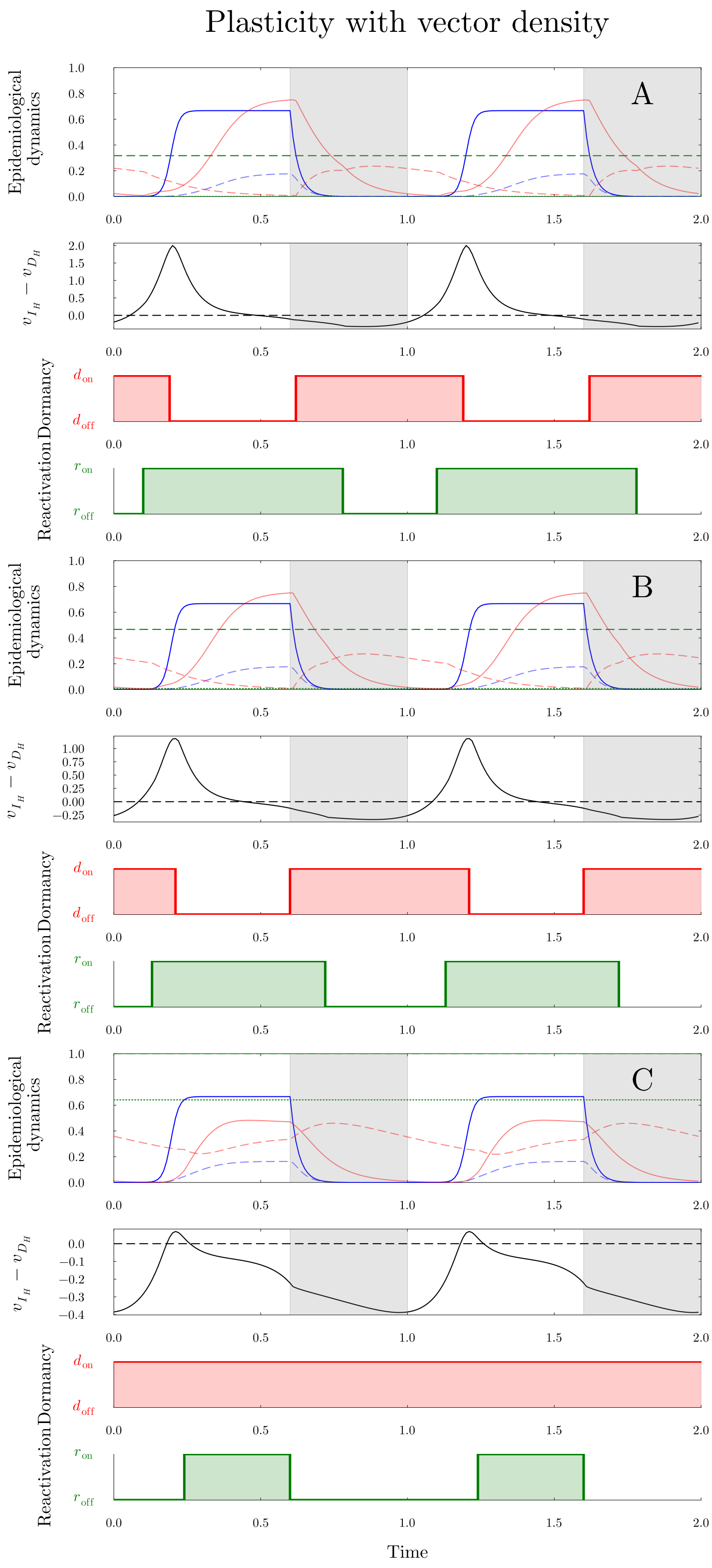


**Figure S7: Evolution of time-varying dormancy and reactivation in a seasonal environment and plasticity is driven by vector density:** in the top plot, the dashed green line is the threshold level $N_V^{d*}$, and the dotted green line is the threshold level $N_V^{r*}$. We vary the cost of dormancy: $c_D = 1$ in (A), $c_D = 0.85$ in (B), $c_D = 0.4$ in (C). Other parameter values are are given in **Table 1**.
